# Cryo-sensitive aggregation triggers NLRP3 inflammasome assembly in cryopyrin-associated periodic syndrome

**DOI:** 10.1101/2021.10.05.463273

**Authors:** Tadayoshi Karasawa, Takanori Komada, Naoya Yamada, Emi Aizawa, Yoshiko Mizushina, Sachiko Watanabe, Chintogtokh Baatarjav, Takayoshi Matsumura, Masafumi Takahashi

## Abstract

Cryopyrin-associated periodic syndrome (CAPS) is an autoinflammatory syndrome caused by mutations of NLRP3, which was originally identified as cryopyrin. Familial cold autoinflammatory syndrome (FCAS), the mildest form of CAPS, is characterized by cold-induced inflammation induced by the overproduction of IL-1β. However, the molecular mechanism of how mutated NLRP3 causes inflammasome activation in CAPS remains unclear. Here, we found that CAPS-associated NLRP3 mutants form cryo-sensitive aggregates that function as a scaffold for inflammasome activation. Cold exposure promoted inflammasome assembly and subsequent IL-1β release triggered by mutated NLRP3. While K^+^ efflux was dispensable, Ca^2+^ was indispensable for mutated NLRP3-mediated inflammasome assembly. Notably, Ca^2+^ influx was induced during mutated NLRP3-mediated inflammasome assembly. Furthermore, caspase-1 inhibition prevented Ca^2+^ influx and inflammasome assembly induced by the mutated NLRP3, suggesting a feed-forward Ca^2+^ influx loop triggered by mutated NLRP3. Thus, the mutated NLRP3 forms cryo-sensitive aggregates to promote inflammasome assembly distinct from canonical NLRP3 inflammasome activation.

## Introduction

The cryopyrin-associated periodic syndromes (CAPS) are a spectrum of rare diseases consisting of three clinically defined autosomal dominant disorders: familial cold autoinflammatory syndrome (FCAS), Muckle-Wells syndrome (MWS), and chronic infantile neurological, cutaneous and articular syndrome (CINCA) (Kuemmerle-Deschner et al., 2017). These three syndromes can be classified according to severity. FCAS is the mildest form of CAPS and is characterized by cold-induced fever and inflammation. MWS is accompanied by systemic amyloidosis and hearing loss. CINCA is the most severe phenotype and is characterized by central nervous system inflammation and bone deformities. Genetic causes of these disorders are gain-of-function mutations in the Nucleotide-binding oligomerization domain, leucine-rich repeat and pyrin domain containing 3 (*NLRP3*) gene, also known as cryopyrin (Brydges et al., 2009; Kuemmerle-Deschner, 2015). The mutated NLRP3 protein causes overproduction of interleukin (IL)-1β, resulting in systemic inflammatory characteristics, such as recurrent fever, rash, conjunctivitis, and arthralgia.

NLRP3 forms a multi-protein molecular complex called “NLRP3 inflammasome” (Schroder and Tschopp, 2010). NLRP3 inflammasome is composed of NLRP3, apoptosis-associated speck-like protein containing a caspase recruitment domain (ASC) which acts as an adaptor protein, and the cysteine proteinase caspase-1, and functions as a scaffold for caspase-1 activation (Schroder and Tschopp, 2010; Karasawa and Takahashi, 2017). The assembly of inflammasome complex promotes oligomerization and auto-processing of caspase-1. The active caspase-1 processes precursors of inflammatory cytokines IL-1β and IL-18 and converts them to their mature forms. Another critical role of caspase-1 is the processing of gasdermin D (GSDMD) (Liu et al., 2016; Broz et al., 2020). The processed amino-terminal domain of GSDMD binds to the plasma membrane and forms pores. Therefore, caspase-1-mediated GSDMD pore induces the release of cytosolic content and subsequent necrotic cell death called pyroptosis.

Although NLRP3 was initially identified as a causative gene of CAPS (Feldmann et al., 2002), the function of NLRP3 had been unclear because CAPS is a rare disease. In 2006, however, Tschopp and his colleagues found that monosodium urate crystals activate NLRP3 inflammasome (Martinon et al., 2006). Since this finding, many studies have clarified the pivotal role of NLRP3 inflammasome in inflammatory responses in both host defense and sterile inflammatory diseases. Other investigators and we have demonstrated the pathophysiological role of NLRP3 inflammasome in cardiovascular and renal diseases (Duewell et al., 2010; Usui et al., 2012, 2015; Komada et al., 2014, 2015).

Despite many findings regarding molecular mechanisms and the pathophysiological role of the NLRP3 inflammasome, the disease mechanisms of CAPS are not fully understood. In particular, although FCAS is characterized by cold exposure-induced recurrent fever and inflammation, the mechanisms by which exposure to cold regulates NLRP3 inflammasome in FCAS remain unclear (Rosengren et al., 2007; Brydges et al., 2013). In the present study, we have found that CAPS-associated NLRP3 mutants form cryo-sensitive aggregates, which function as scaffolds for NLRP3 inflammasome assembly. The aggregation of the mutated NLRP3 is sensitive to Ca^2+^. Therefore, mutated NLRP3 triggers inflammasome assembly driven by Ca^2+^ influx-mediated feed-forward regulation.

## Results

### CAPS-associated NLRP3 mutants form cryo-sensitive foci

To investigate the pathophysiological role of CAPS-associated NLRP3 mutants, we generated cell lines expressing fusion proteins of NLRP3 mutants and a green monomeric protein, mNeonGreen (Figure 1–figure supplement1A and B). We found that FCAS-associated NLRP3-L353P and CINCA-associated NLRP3-D303N formed foci without any stimulation, while wild type (WT)-NLRP3 is expressed diffusely (Figure 1A). On the other hand, ASC-GFP forms a single speck per cell. To analyze the localization of NLRP3 during NLRP3 inflammasome activation without being affected by ASC, we generated *ASC KO* THP-1 cells (*ASC KO/EF-1-NLRP3-mNeonGreen*-THP-1). NLRP3-D303N formed foci in *ASC KO* THP-1 cells, whereas WT-NLRP3 did not form foci upon stimulation by the NLRP3 activator nigericin, indicating that the foci are distinct from canonical inflammasome assembly (Figure 1B and C). To assess whether the foci formation is cryo-sensitive, the transduced cells were exposed to cold temperature (32°C) for 24 h. A considerable number of foci were detected only in D303N mutant-expressing cells under normal temperature (37°C). However, the number of foci was increased both in FCAS-associated L353P and CINCA-associated D303N mutant-expressing cells under cold exposure (Figure 1D–I). The number of foci formed was weakly associated with expression levels of NLRP3 as indicated by fluorescence of mNeonGreen. In contrast, speck formation by ASC-GFP was not affected by cold exposure (Figure 1J and Figure 1–figure supplement1C). These results suggest that CAPS-associated NLRP3 mutants form cryosensitive foci consistent with disease severity and characteristics.

**Figure 1.**
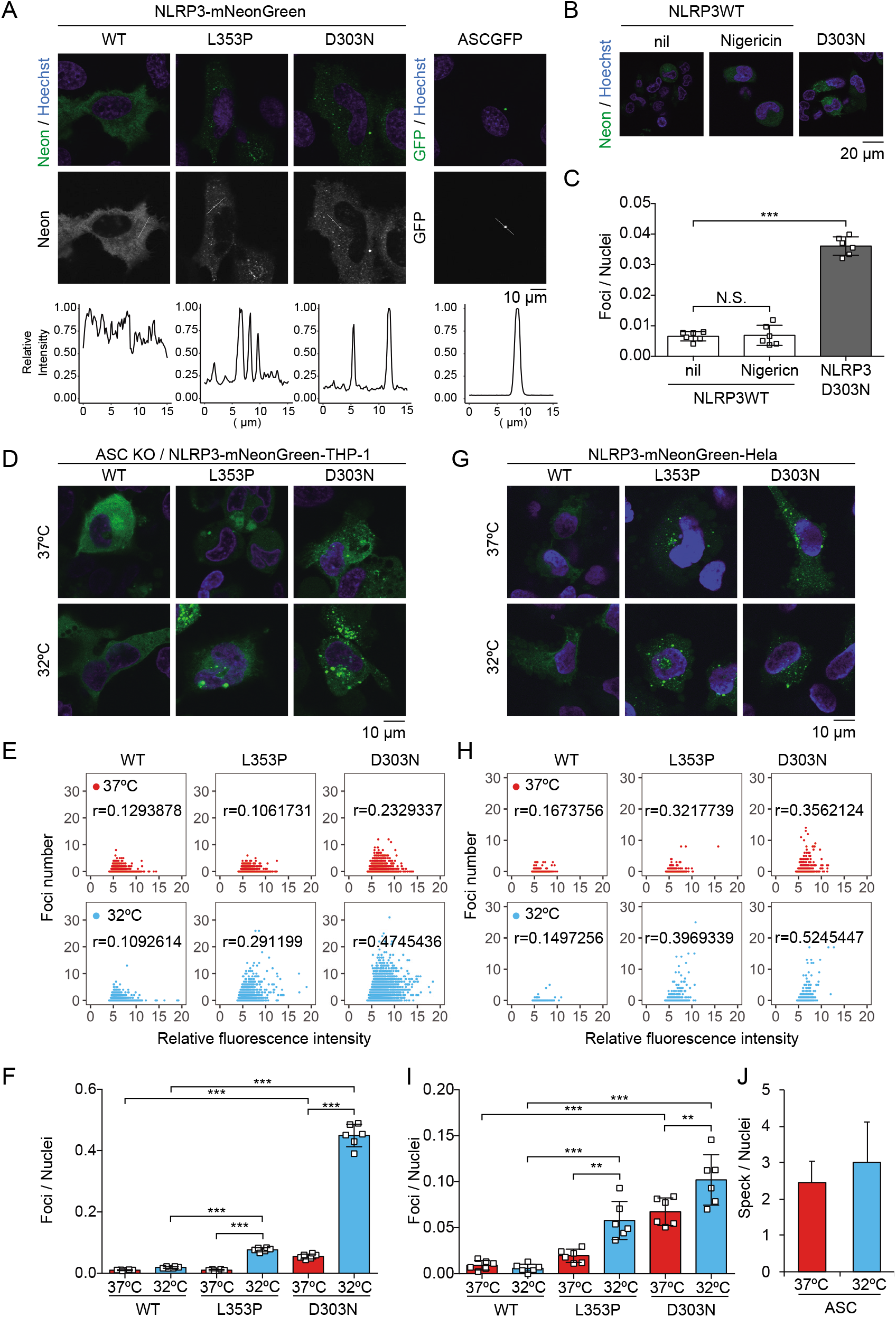
CAPS-associated NLRP3 mutants form cryo-sensitive foci. (A) *EF1-NLRP3-WT-, NLRP3-L353P*-, or *NLRP3-D303N-mNeonGreen*-Hela cells or *EF1-ASCGFP*-Hela cells were analyzed by confocal microscopy. Line profiles of foci or specks in the images were analyzed. (B and C) *ASC KO/EF1-NLRP3-WT*-, or *NLRP3-D303N-mNeonGreen*-THP-1 cells were differentiated with 200 nM PMA for 24 h and then treated with nigericin for 6 h. (B) Representative images by confocal microscopy. (C) The number of foci was counted by high content analysis. (D–F) Differentiated *ASC KO/EF1-NLRP3-WT-, NLRP3-L353P*-, or *NLRP3-D303N-mNeonGreen*-THP-1 cells were cultured at 37°C or 32°C for 24 h. (G–I) *EF1-NLRP3-WT-, NLRP3-L353P*-, or *NLRP3-D303N-mNeonGreen*-HeLa cells were cultured at 37°C or 32°C for 24 h. (D and G) Representative images by confocal microscopy. (E, F, H, I) The number of foci and the fluorescence intensity of the cells were analyzed by high content analysis. Pearson correlation coefficients are shown. (J) *EF1-ASC-GFP*-HeLa cells were cultured at 37°C or 32°C for 24 h. The number of nuclei and speck was counted. (C, F, I, J) Data are expressed as the mean ± SD. \**P* < 0.05, \*\**P* < 0.01, \*\*\**P* < 0.005 as determined by two-way ANOVA with a post hoc test. Data are representative of three independent experiments.

**Figure 1 – figure supplement 1.**
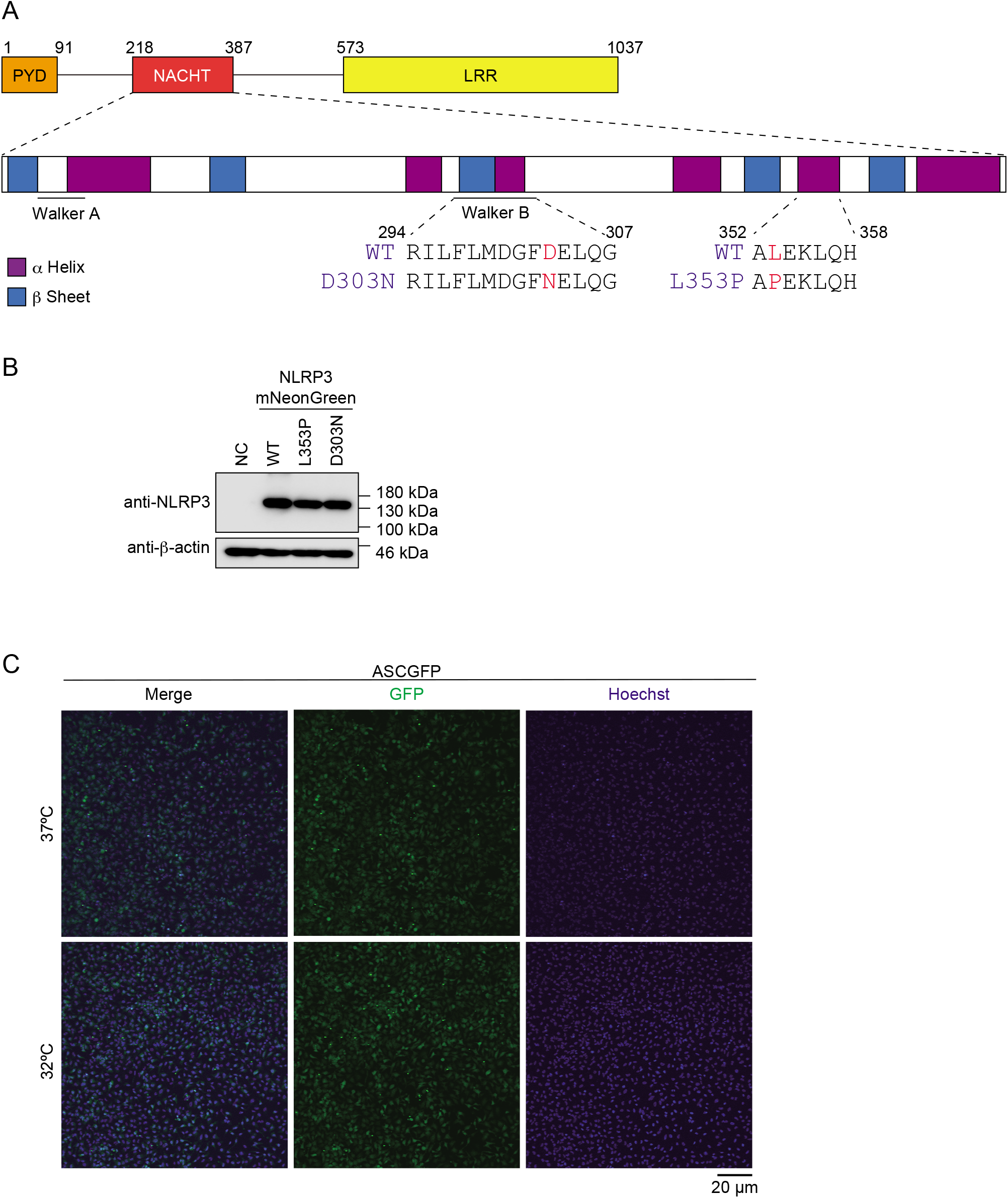
Expression of NLRP3-mNeonGreen and ASC-GFP. (A) Amino acid sequence of CINCA-associated D303N mutant and FCAS-associated L353P mutant. (B)Lysates of *EF1-NLRP3-WT-, NLRP3-L353P*-, or *NLRP3-D303N-mNeonGreen* HeLa cells were analyzed by western blot. (C) *EF1-ASC-GFP*-HeLa cells were cultured at 37°C or 32°C for 24 h. Representative image of confocal microscopy.

### CAPS-associated NLRP3 mutants form aggregates

Since NLRP3 has a PYD scaffold domain (Figure 2–figure supplement 1A), a common feature of molecules that form aggregates or liquid-liquid-phase separation (LLPS), we hypothesized that CAPS-associated NLRP3 mutants form aggregates or LLPS (Alberti et al., 2019). To test this hypothesis, we performed fluorescence recovery after photobleaching (FRAP) analysis. After induction of foci formation by cold exposure, some of the NLRP3-L353P- and D303N-mNeonGreen-foci were bleached. The fluorescence in the bleached area was not recovered, indicating that the foci are aggregates (Figure 2A and B, Figure 2–figure supplement 2A and B). The fully bleached area of NLRP3-L353P-mNeonGreen-foci was also not recovered (Figure 2–figure supplement 2C and D). Similar results are obtained from FRAP analysis of ASC speck, initially reported to be aggregates (Masumoto et al., 1999) (Figure 2C and D). In both NLRP3-L353P foci and ASC speck, the exchange of protein between bleached and unbleached area was not detected (Figure 2E and F). Furthermore, 1,6-hexanediol, an LLPS inhibitor, did not affect foci formation of NLRP3-mNeonGreen (Figure 2–figure supplement 2E). These results suggest that foci formed by CAPS-associated NLRP3 mutants are aggregates.

**Figure 2.**
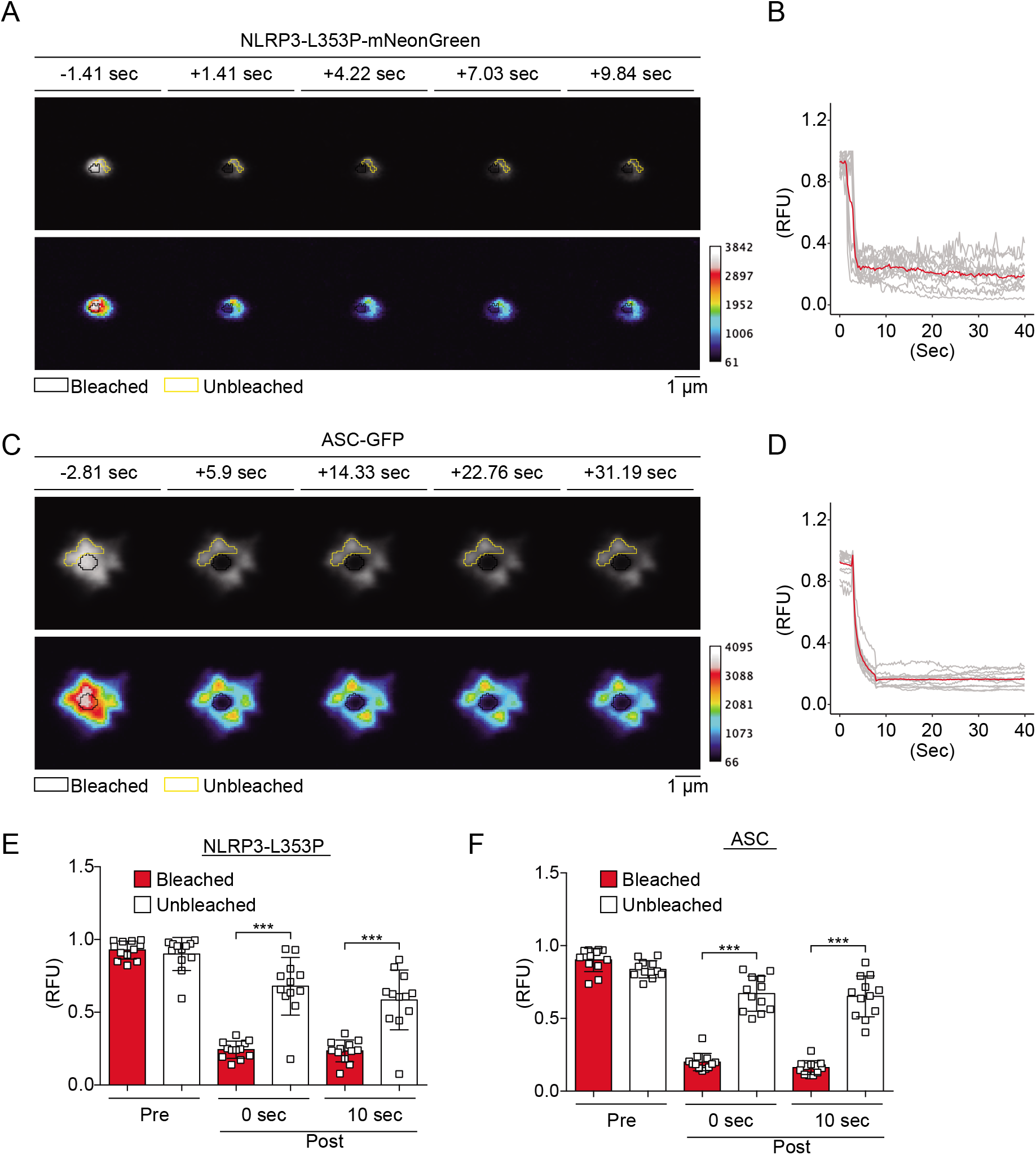
CAPS-associated NLRP3 mutants form aggregates. (A–F) *EF1-NLRP3-L353P-mNeonGreen*- or *EF1-ASC-GFP*-HeLa cells were cultured at 32°C for 24 h. Foci or specks in the cells were analyzed by FRAP. Representative images of (A) foci formed by *NLRP3-L353P-mNeonGreen* or (C) specks formed by *ASC-GFP* before and after photobleaching. The bleached and unbleached areas are shown in black lines and yellow lines, respectively. Plots of relative fluorescence units during photobleaching of (B and E) *NLRP3-L353Pm-NeonGreen* (n = 12) and (D and F) ASC specks (n = 12). (B and D) The red line represents mean values and the gray lines represent each measurement. (E and F) Data are expressed as the mean ± SD. \**P* < 0.05, \*\**P* < 0.01, \*\*\**P* < 0.005 as determined by two-way ANOVA with a post hoc test. Data are from three independent live-cell imaging.

### Aggregates formed by CAPS-associated NLRP3 mutants are the scaffold for inflammasome activation

Next, we investigated whether aggregates formed by mutated NLRP3 function as a scaffold for inflammasome assembly and trigger subsequent IL-1β release. In order to analyze colocalization of NLRP3 and ASC, we developed THP-1 cells harboring two reporters; *TRE-NLRP3-mNeonGreen* and *EF1-ASC-BFP*. After induction of NLRP3-L353P-mNeonGreen by doxycycline (DOX), ASC-speck was colocalized with the NLRP3 mutant-formed aggregate. (Figure 3A–C). To exclude the possibility that NLRP3 mutant aggregation is due to its fluorescent tag, the cells expressing NLRP3 mutants under TET-ON promoter were developed (Figure 3–figure supplement 1A and B). In accordance with the cryo-sensitive formation of aggregates by NLRP3-L353P, insoluble complex formation and oligomerization of ASC induced by NLRP3-L353P were enhanced by cold exposure (Figure 3D and E). In contrast, the ASC-oligomerization induced by nigericin was attenuated under cold exposure (Figure 3F). ASC-speck formation was further assessed by fusion protein of ASC-GFP reporter. Similarly, NLRP3-L353P-induced ASC speck formation was increased under cold exposure (Figure 3G). Moreover, cold exposure enhanced IL-1β release induced by the NLRP3-L353P mutant, whereas nigericin- and nanosilica-induced IL-1β release was restrained under cold exposure (Figure 3H and Figure 3–figure supplement 1C). Cold exposure also enhanced IL-1β release in CINCA-associated NLRP3-D303N-expressing cells (Figure 3–figure supplement 1D). In contrast, WT-NLRP3 failed to induce IL-1β release (Figure 3–figure supplement 1E and F). These results demonstrate that cryo-sensitive aggregates formed by CAPS-associated NLRP3 mutants function as a scaffold for inflammasome activation and induce subsequent IL-1β release.

**Figure 2 – figure supplement 1.**
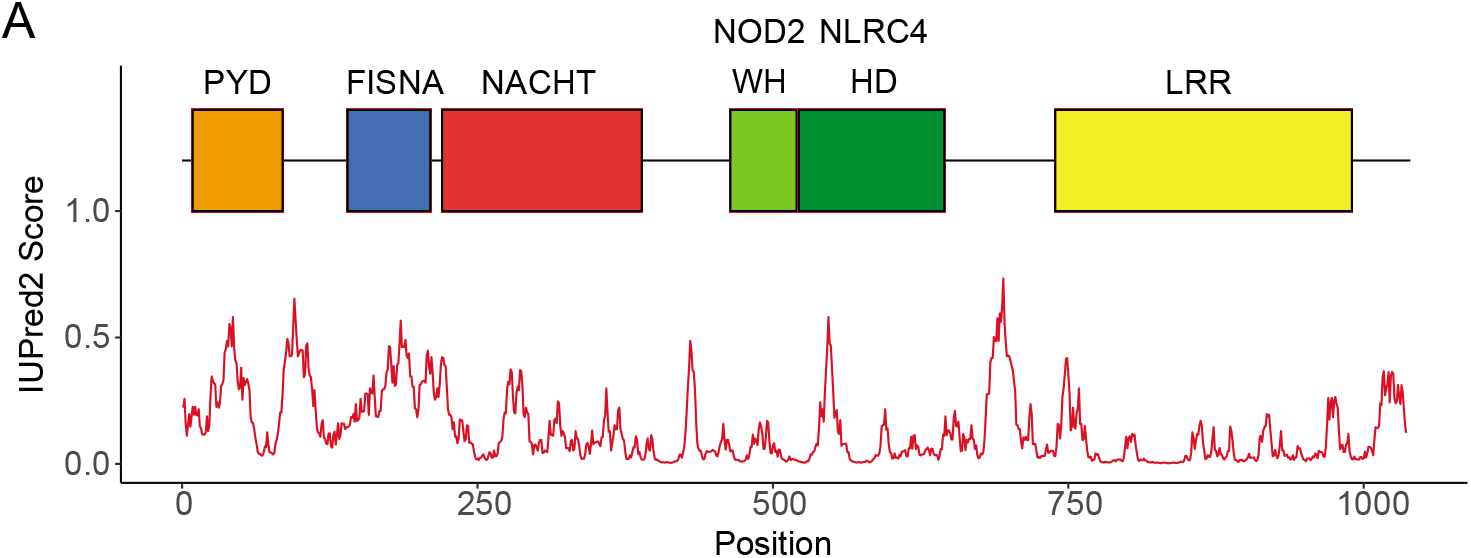
Domains composing NLRP3. (A) IUPred score of NLRP3 and domains in human NLRP3.

**Figure 2 – figure supplement 2.**
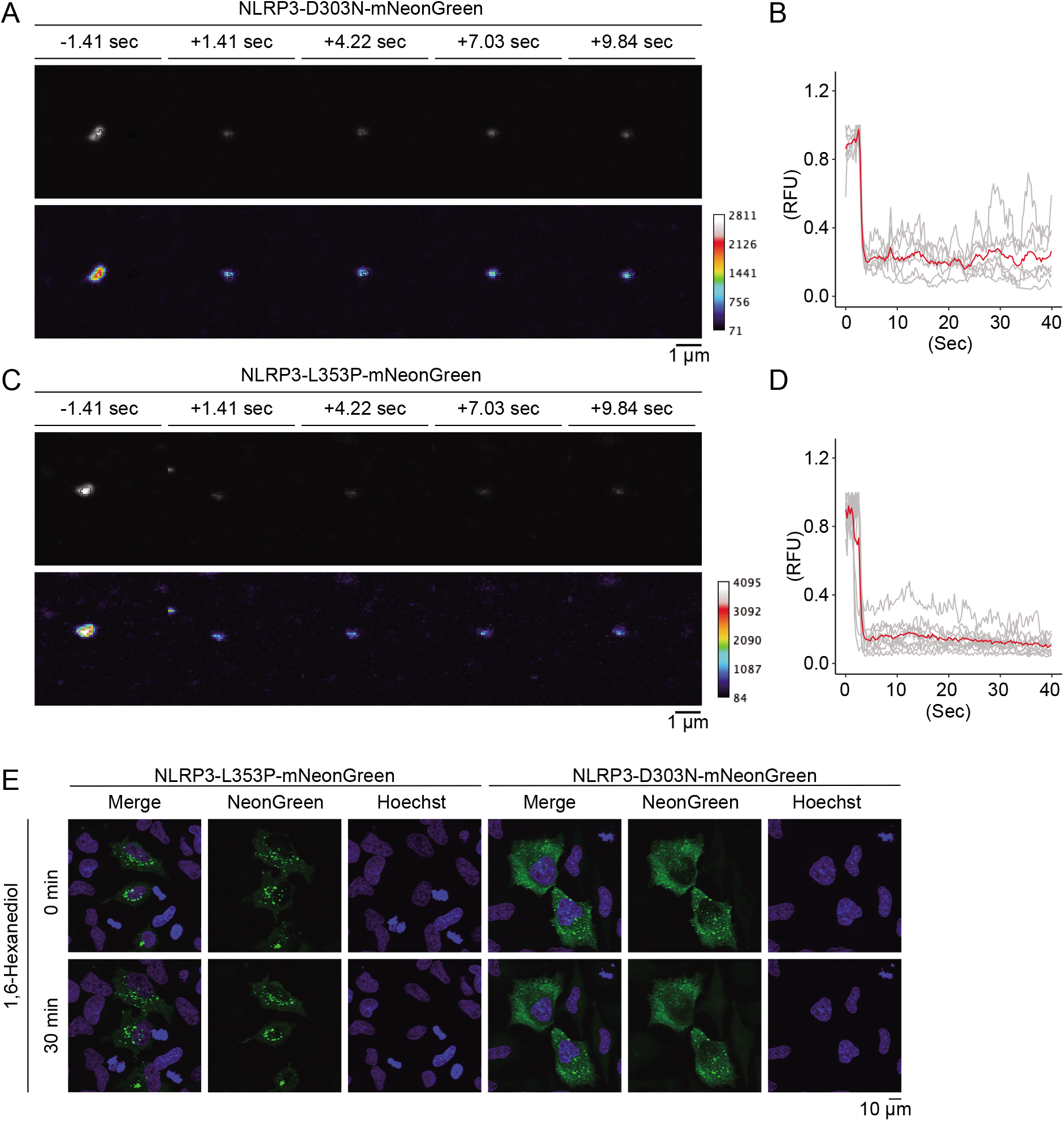
CAPS-associated NLRP3 mutants form aggregates. (A–D) *EF1-NLRP3-L353P*-, or *NLRP3-D303N-mNeonGreen* HeLa cells were cultured at 32°C for 24 h. Foci in the cells were analyzed by fluorescence recovery after photobleaching. Representative images of foci formed (A) by *NLRP3-D303N-mNeonGreen* or (C) by *NLRP3-L353P-mNeonGreen* before and after photobleaching. The bleached area and unbleached area are shown in black lines and yellow lines, respectively. Plot of relative fluorescence unit during photobleaching of (B) *NLRP3-D303N-mNeonGreen* (n = 7) and (D) *NLRP3-L353P-mNeonGreen* (n = 10). The red line represents the mean value and the gray lines represent each measurement. Data are from two or three independent live-cell imaging. (E) *EF1-NLRP3-L353P*-, or *NLRP3-D303N-mNeonGreen* HeLa cells were treated with 5% 1,6-Hexanediol for 30 min. Representative images of live-cell imaging.

**Figure 3.**
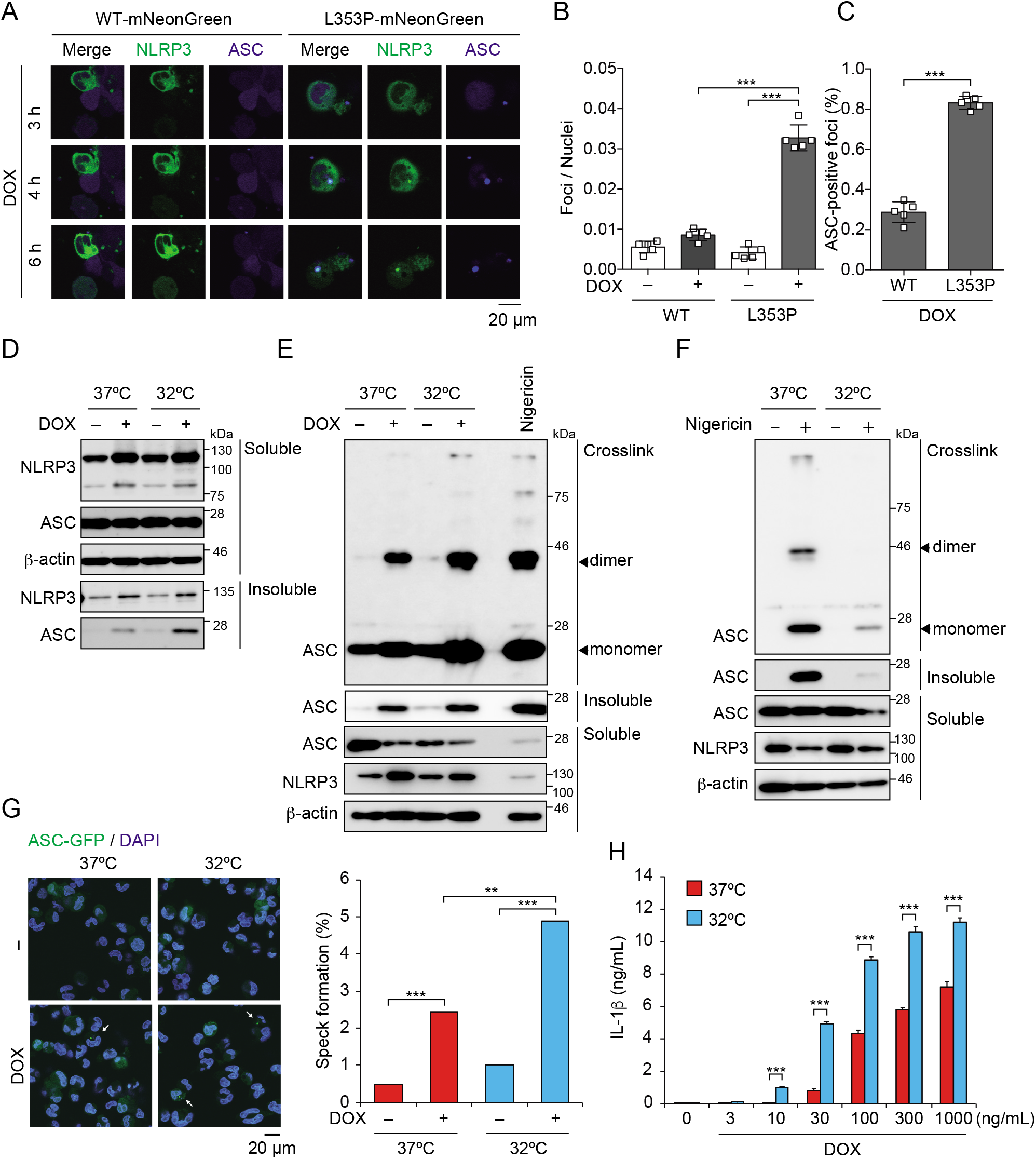
Aggregates formed by CAPS-associated NLRP3 mutant are the scaffold for inflammasome activation. (A–C) *EF1-ASC-BFP/TRE-NLRP3-WT* or *L353P-mNeonGreen* THP-1 cells were treated with DOX. (A) Localization of ASC and NLRP3 was analyzed by confocal microscopy. (B) The number of foci was counted. (C) The ASC speck number in NLRP3 foci was analyzed. (D and E) *TRE-NLRP3-L353P*-THP-1 cells were differentiated with PMA for 24 h and then treated with DOX (30 ng/mL) at 37°C or 32°C for 6 h. (D) Triton X-soluble and -insoluble fractions were analyzed by Western blot. (E) Oligomerized ASC in Triton X-insoluble fractions was crosslinked with BS3 and analyzed by Western blot. (F) Differentiated *TRE-NLRP3-L353P*-THP-1 cells were treated with nigericin at 37°C or 32°C for 6 h. Triton X-insoluble fractions were crosslinked with BS3 and analyzed by Western blot. (G) *EF-1-ASC-GFP/TRE-NLRP3-L353P*-THP-1 cells were differentiated with PMA for 24 h and then treated with DOX (30 ng/mL) at 37°C or 32°C for 6 h. Representative images by confocal microscopy and the number of nuclei and specks were counted. (H) Differentiated *TRE-NLRP3-L353P*-THP-1 cells were treated with DOX at 37°C or 32°C for 6 h. The IL-1β levels in the supernatants were assessed by ELISA (n = 3). (B, C and H) Data are expressed as the mean ± SD. \**P* < 0.05, \*\**P* < 0.01, \*\*\**P* < 0.005 as determined by (B, C, and H) two-way ANOVA with a post hoc test or (G) Fisher’ s exact test with the Holm correction. Data are representative of two or three independent experiments.

### Ca^2+^ is required for NLRP3 mutant-mediated inflammasome assembly

To elucidate the regulatory mechanisms of NLRP3 mutant-mediated inflammasome activation under cold exposure, the upstream pathways of canonical inflammasome including K^+^ efflux, lysosomal destabilization, mitochondrial ROS generation, and Ca^2+^ mobilization were explored. Unexpectedly, inhibition of K^+^ efflux failed to prevent NLRP3-L353P mutant-induced IL-1β release, while it inhibited nigericin-induced IL-1β release (Figure 4–figure supplement 1A). Similarly, deficiency of NEK7, an essential component K^+^ efflux-mediated NLRP3 inflammasome, failed to inhibit NLRP3-L353P mutant-mediated IL-1β release, although it inhibited nigericin-induced IL-1β release (Figure 4–figure supplement 1B–D). Further, inhibition of lysosomal or mitochondrial ROS pathway did not suppress NLRP3-L353P mutant-induced IL-1β release (Figure 4–figure supplement 2A and B). On the other hand, EGTA, a chelator of Ca^2+^, inhibited NLRP3-L353P mutant-induced IL-1β release (Figure 4A, Figure 4–figure supplement 2A and C). Moreover, decreased IL-1β release by NLRP3-L353P mutant under Ca^2+^-depleted conditions was restored by Ca^2+^ supplementation at 32°C (Figure 4B). Next, we assessed whether deprivation or supplementation of Ca^2+^ alters ASC oligomerization. Ca^2+^ deprivation by EGTA attenuated NLRP3-L353P-induced ASC oligomerization, whereas reduced ASC oligomerization in Ca^2+^-depleted conditions was restored by Ca^2+^ supplementation (Figure 4C and Figure 4–figure supplement 2D). The requirement of Ca^2+^ for inflammasome assembly was also confirmed by the use of ASC-GFP reporter cells (Figure 4D and Figure 4–figure supplement 2E). The effect of Ca^2+^ on NLRP3 aggregation was analyzed using NLRP3-mNeonGreen reporter cells. Cryo-sensitive aggregation of NLRP3-L353P was decreased by Ca^2+^-depletion (Figure 4E and F). A similar dependency on Ca^2+^ was also detected in the CINCA-associated NLRP3-D303N mutant (Figure 4–figure supplement 3A). Moreover, DOX-induced NLRP3-L353P aggregation and ASC speck formation were attenuated by Ca^2+^-depletion (Figure 4G and H). These results suggest that Ca^2+^ is an indispensable regulator of the aggregation of CAPS-associated NLRP3 mutants and subsequent activation of NLRP3 inflammasome.

**Figure 3 – figure supplement 1.**
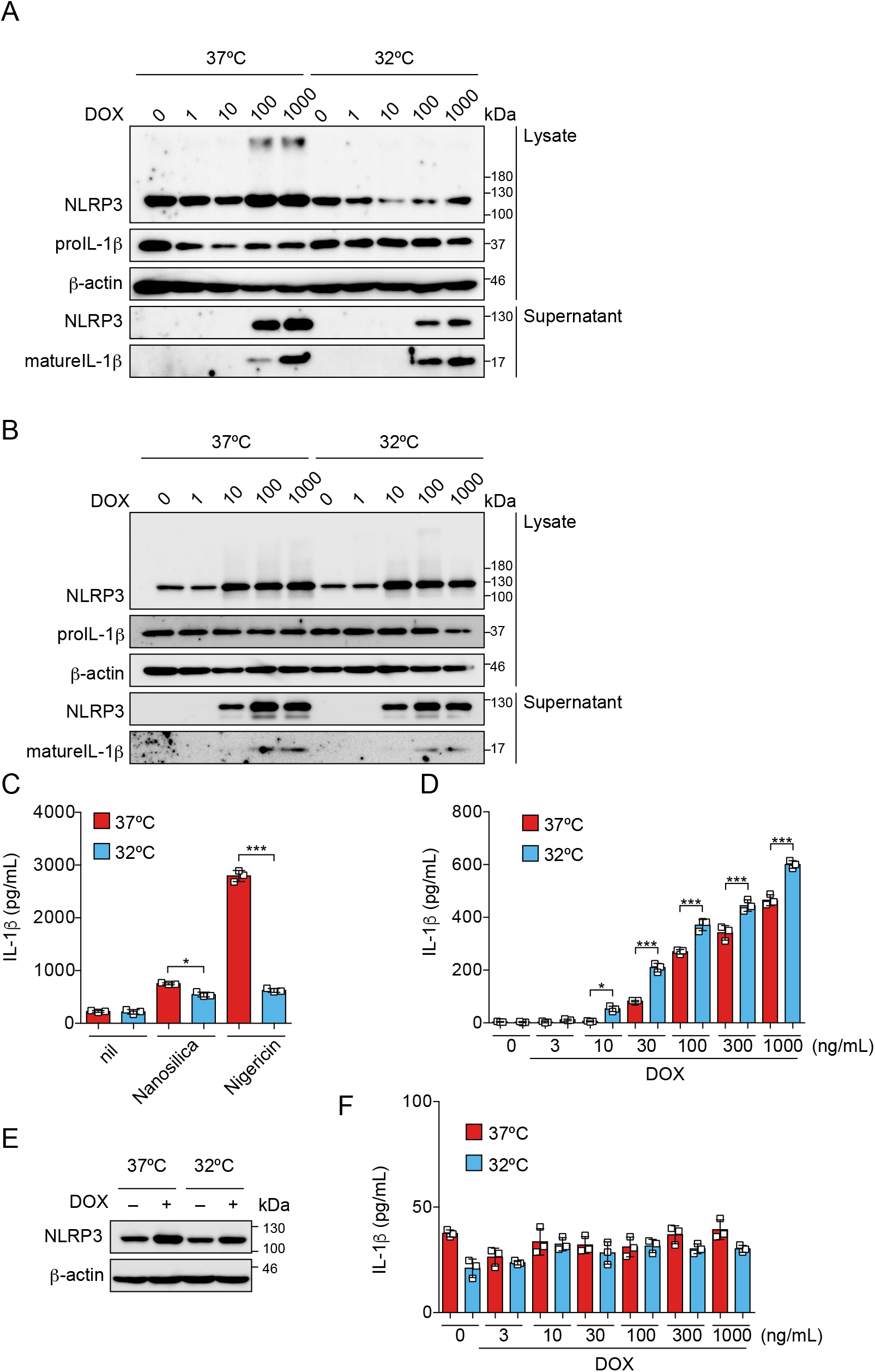
Cold exposure enhances IL-1β release in NLRP3 FCAS mutant-expressing cells. (A) *TRE-NLRP3-L353P*-or (B, D) *NLRP3-D303N*- or (E, F) *NLRP3WT*-THP-1 cells were differentiated with PMA for 24 h and then treated with DOX at 37°C or 32°C for 6 h. (C) Differentiated TRE-NLRP3-L353P-THP-1 cells were treated with 5 μM nigericin or nanosilica (30 μg/mL) at 37°C or 32°C for 6 h. (A, B, E) Lysates and supernatants were analyzed by western blot. (C, D, F) The levels of IL-1β in the supernatants were assessed by ELISA (n = 3). Data are expressed as the mean ± SD. \**P* < 0.05, \*\**P* < 0.01, \*\*\**P* < 0.005 as determined by two-way ANOVA with a post hoc test. Data are representative of two independent experiments.

**Figure 4.**
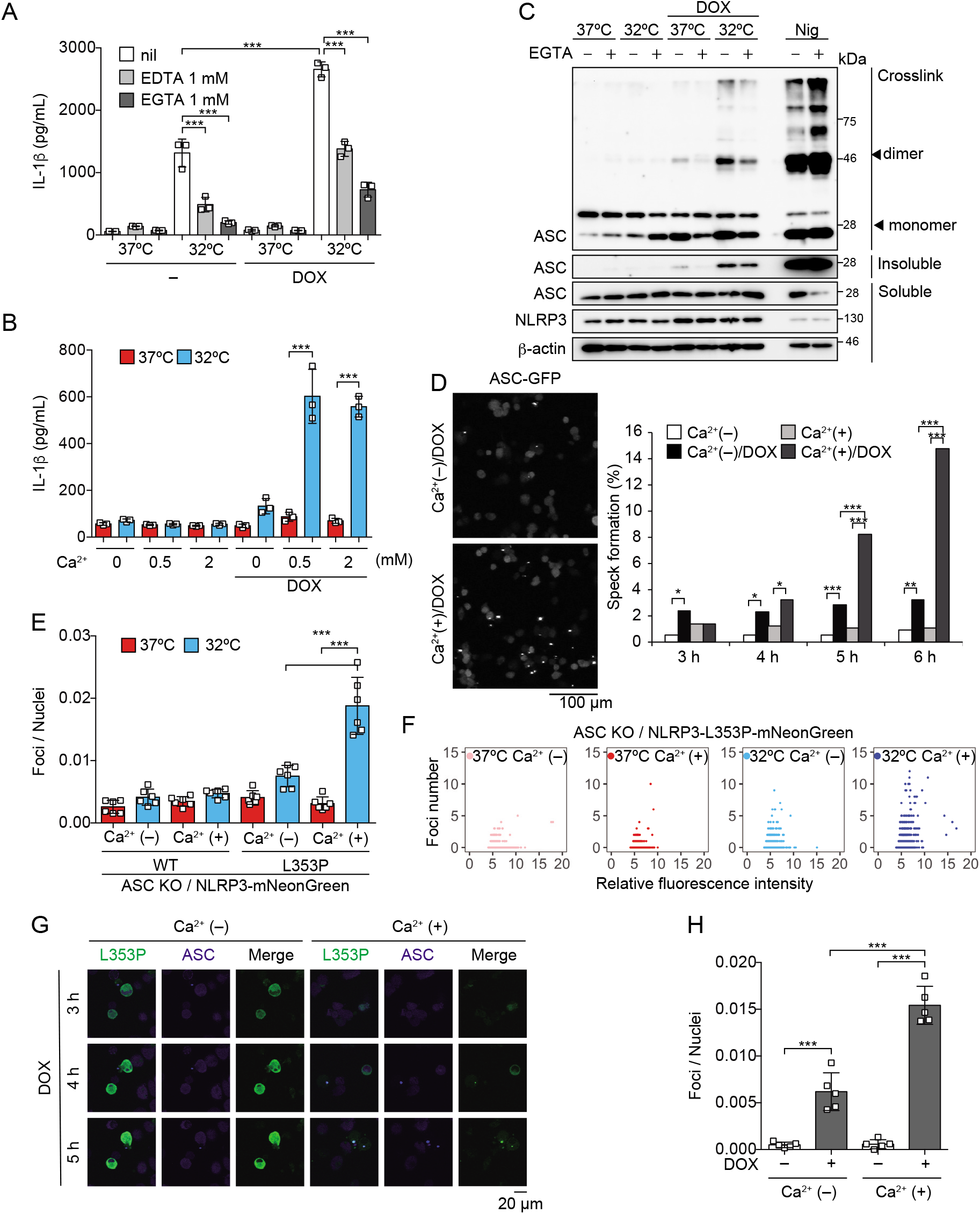
Ca^2+^ is required for CAPS-associated NLRP3 mutant-mediated inflammasome assembly. (A–C) Differentiated *TRE-NLRP3-L353P*-THP-1 cells were pretreated with indicated conditions, and then treated with DOX (30 ng/mL) at 37°C or 32°C for 6 h. (A and B) The IL-1β levels in the supernatants were assessed by ELISA (n = 3). (C) Oligomerized ASC in Triton X-insoluble fractions was crosslinked with BS3 and analyzed by Western blot. (D) *EF1-ASC-GFP/TRE-NLRP3-L353P*-THP-1 cells were pretreated with Ca^2+^-depleted or -supplemented media and then treated with DOX (30 ng/mL) at 37°C. ASC speck formation was analyzed by confocal microscopy. (E and F) *EF1-NLRP3-WT*- or *L353P-mNeonGreen*-THP-1 cells were cultured at 37°C or 32°C for 24 h in Ca^2+^-depleted or -supplemented media. The number of foci and fluorescent intensity were analyzed by high content analysis. (G and H) Differentiated *EF1-ASC-BFP/TRE-NLRP3-L353P-mNeonGreen*-THP-1 cells were pretreated with Ca^2+^-depleted or -supplemented media and then treated with DOX (30 ng/mL) at 37°C. (G) Representative images by confocal microscopy (H) The number of foci was analyzed by high content analysis. (A, B, E, and H) Data are expressed as the mean ± SD. \**P* < 0.05, \*\**P* < 0.01, \*\*\**P* < 0.005 as determined by (A, B, E, and H) two-way ANOVA with a post hoc test or (D) Fisher’ s exact test with the Holm correction. Data are representative of two or three independent experiments.

### Ca^2+^ influx is provoked during mutated NLRP3-mediated inflammasome assembly

To further investigate the role of Ca^2+^ in mutated NLRP3-mediated inflammasome activation, we monitored changes in Ca^2+^ levels using Fluo-8, a fluorescent Ca^2+^ indicator. After DOX-mediated induction of NLRP3-L353P, Ca^2+^ increase was clearly detected (Figure 5A–C). This increased intracellular Ca^2+^ is due to influx because Ca^2+^ increase was not observed in the absence of extracellular Ca^2+^ (Figure 5D–F, Video 1). The increased Ca^2+^ levels were not provoked by membrane rupture because Ca^2+^ influx occurred prior to the release of cytosolic content as indicated by Kusabira orange or membrane permeabilization as indicated by SYTOX (Figure 5–figure supplement 1A–D). Notably, inflammasome assembly monitored by ASC-BFP and Ca^2+^ increase occurred coincidentally (Figure 5G–I). The size of ASC speck increased with the increase in Ca^2+^. These results indicate that Ca^2+^ influx occurs during inflammasome activation induced by NLRP3 mutants.

**Figure 4 – figure supplement 1.**
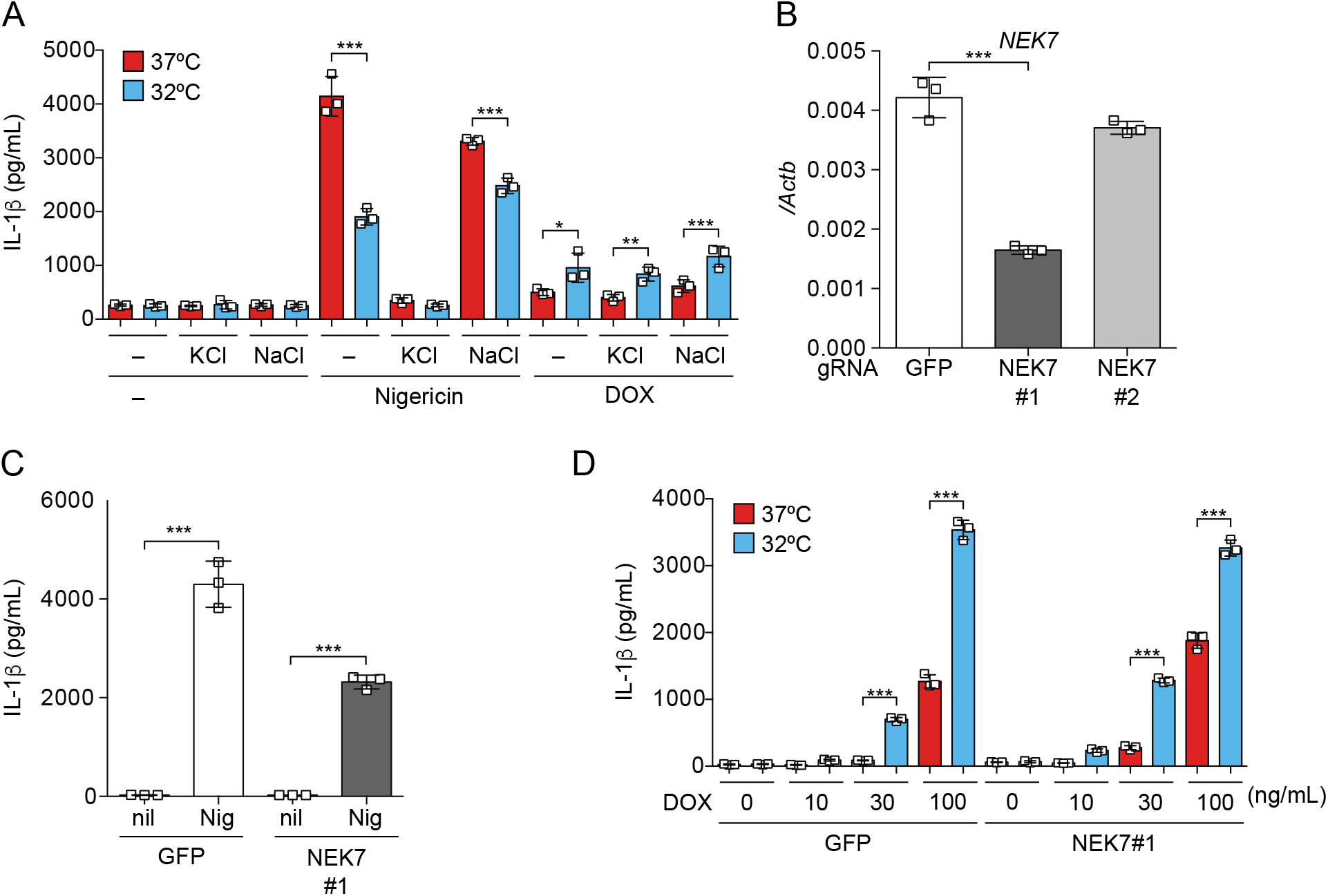
K^+^ efflux is dispensable for inflammasome activation induced by NLRP3 FCAS-mutant. (A) Differentiated *TRE-NLRP3-L353P*-THP-1 cells were pretreated with the indicated dose of KCl or NaCl and then treated with DOX (30 ng/mL) at 37°C or 32°C for 6 h. (B–D) NEK7-mutated *TRE-NLRP3-L353P*-THP-1 cells were differentiated with PMA for 24 h. (B) mRNA expression of NEK7 was analyzed by qPCR (n=3). The differentiated NEK7-mutated NLRP3-L353P-THP-1 cells were treated with (C) nigericin or (D) DOX (30 ng/mL) at 37°C or 32°C for 6 h. The levels of IL-1β in the supernatants were assessed by ELISA (n=3). The levels of IL-1β in the supernatants were assessed by ELISA (n=3). (A–D) Data are expressed as the mean ± SD. \**P* < 0.05, \*\**P* < 0.01, \*\*\**P* < 0.005 as determined by two-way ANOVA with a post hoc test. Data are representative of two independent experiments.

**Figure 4 – figure supplement 2.**
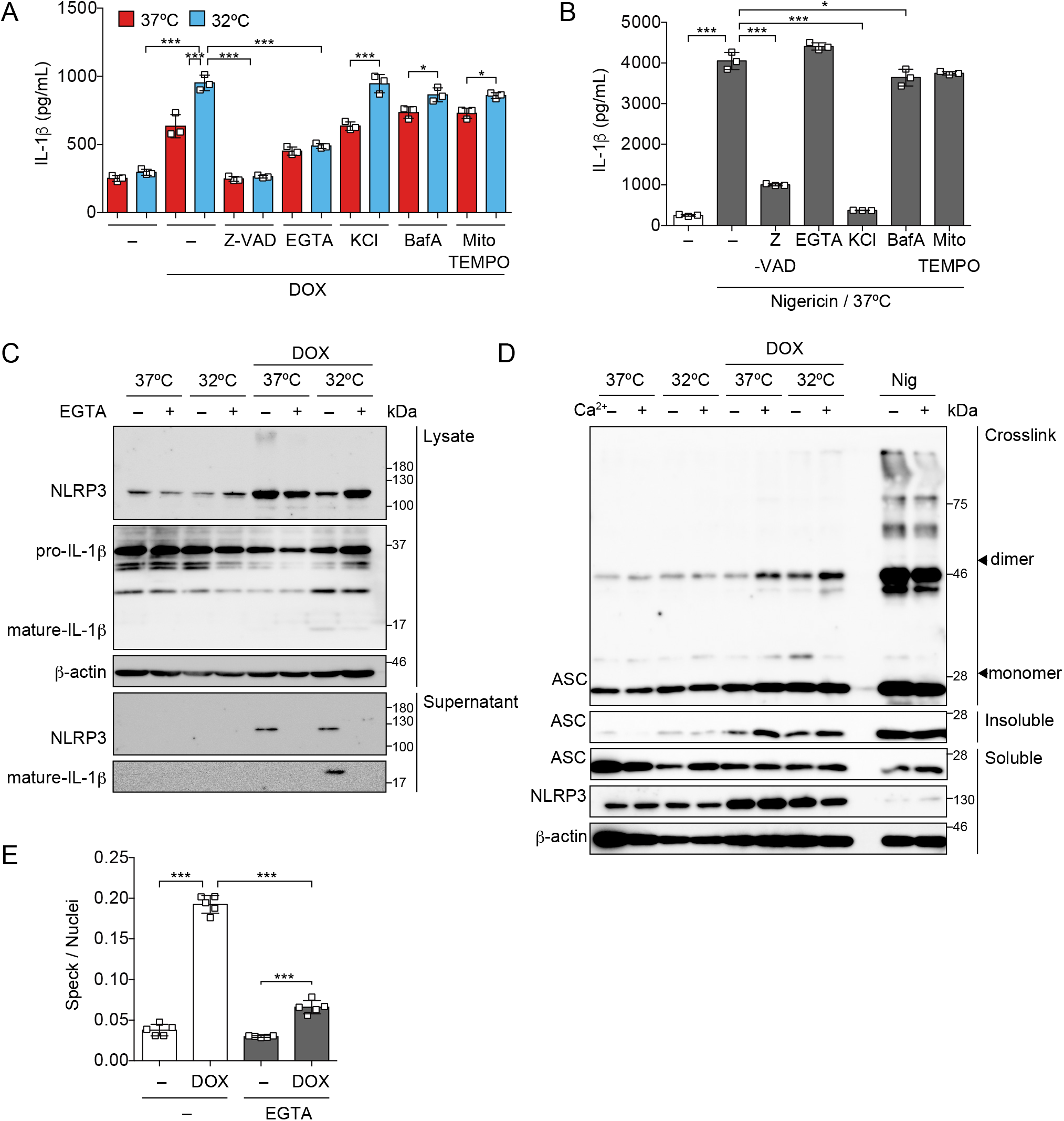
Ca^2+^ is necessary for inflammasome activation induced by mutated NLRP3. (A and B) Differentiated *TRE-NLRP3-L353P*-THP-1 cells were pretreated with the indicated inhibitors and then treated with DOX (30 ng/mL) or nigericin (5 μM) at 37°C or 32°C for 6 h. The levels of IL-1β in the supernatants were assessed by ELISA (n=3). (C) Differentiated *TRE-NLRP3-L353P*-THP-1 cells were pretreated with EGTA and then treated with DOX (30 ng/mL) at 37°C or 32°C for 6 h. Lysates and supernatants were analyzed by western blot. (D) Differentiated *TRE-NLRP3-L353P*-THP-1 cells were pretreated with Ca^2+^-depleted or -supplemented media and then treated with DOX (30 ng/mL) at 37°C or 32°C for 6 h. Oligomerized ASC in Triton X-insoluble fractions was crosslinked by BS3 and analyzed by western blot. (E) *EF1-ASC-GFP/TRE-NLRP3-L353P*-THP-1 cells were pretreated with EGTA and then treated with DOX (30 ng/mL) at 37°C. The formation of ASC-speck was analyzed by high content analysis. (A, B and E) Data are expressed as the mean ± SD. \**P* < 0.05, \*\**P* < 0.01, \*\*\**P* < 0.005 as determined by two-way ANOVA with a post hoc test. Data are representative of two or three independent experiments.

**Figure 4 – figure supplement 3.**
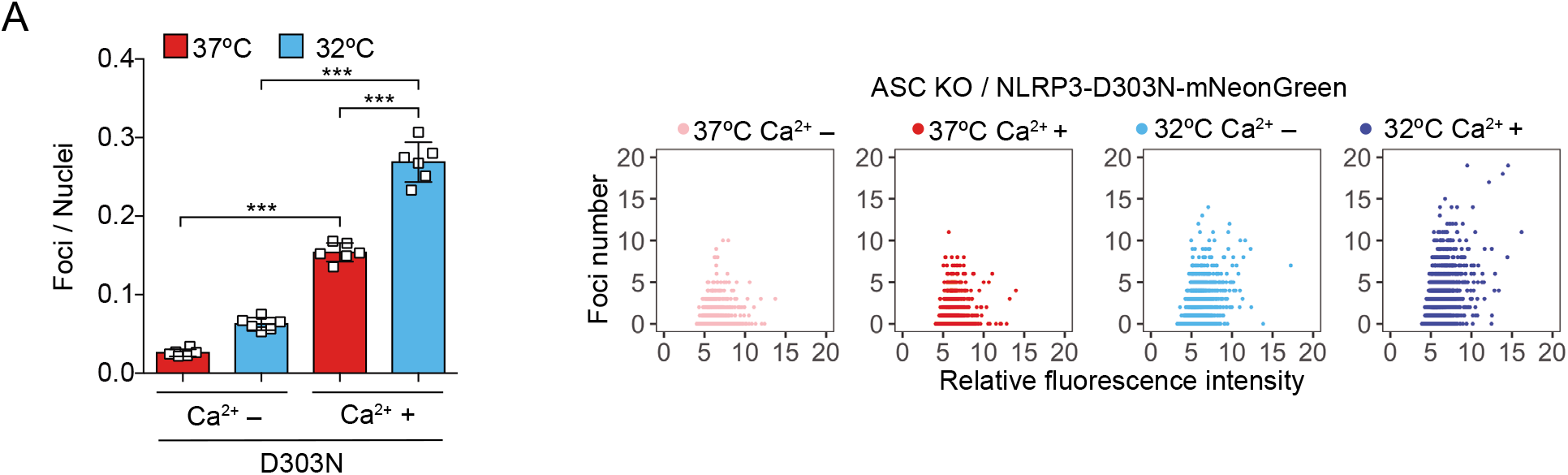
Ca^2+^ is necessary for aggregation of NLRP3-D303N-mutant. (A) *EF1-NLRP3-D303N-mNeonGreen*-THP-1 cells were cultured at 37°C or 32°C for 24 h in Ca^2+^-depleted or -supplemented media. The number of foci and fluorescent intensity were analyzed by high content analysis. Data are expressed as the mean ± SD. \*\*\**P* < 0.005 as determined by two-way ANOVA with a post hoc test. Data are representative of three independent experiments.

**Figure 5.**
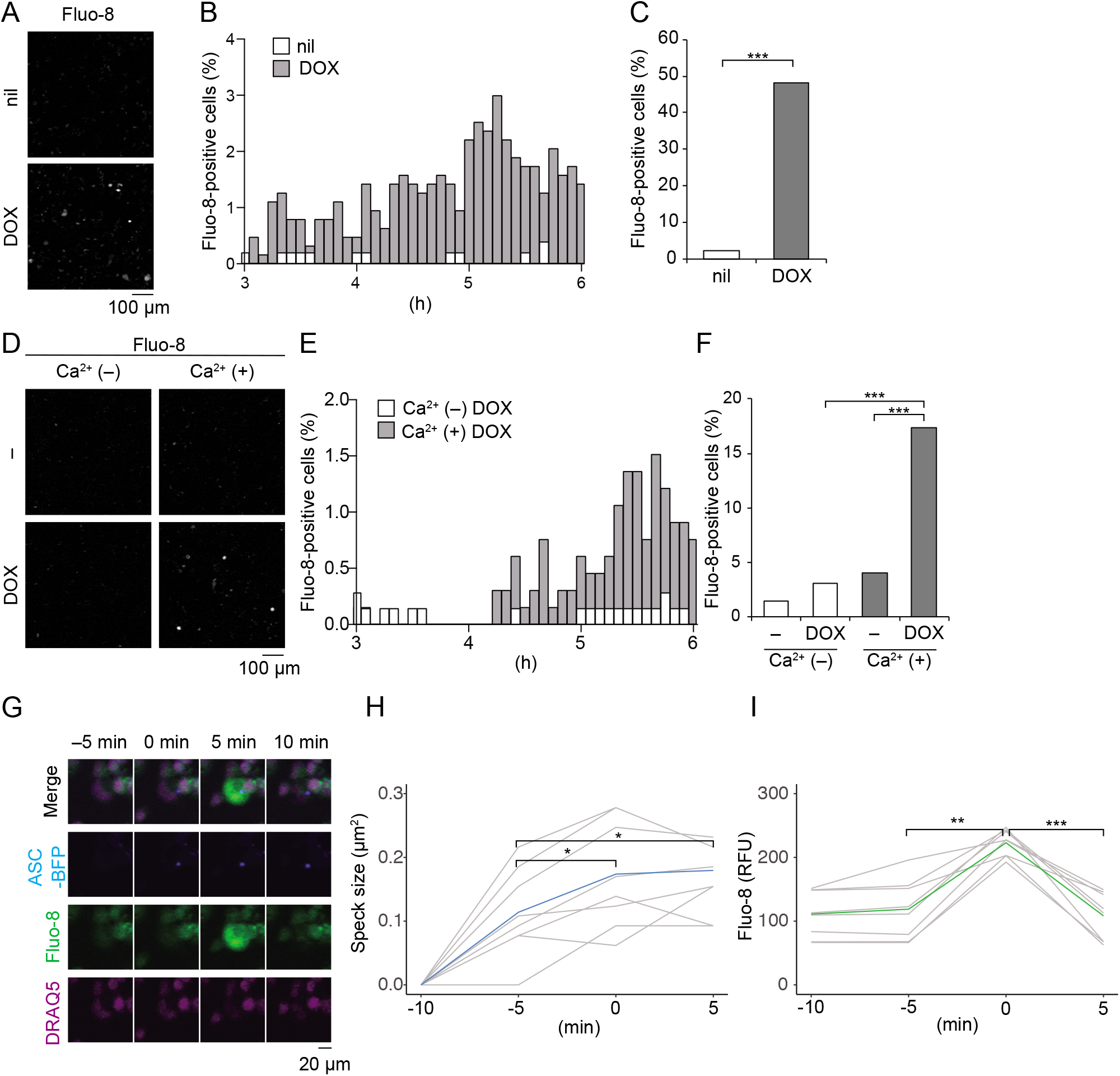
Ca^2+^ influx is provoked during mutated NLRP3-mediated inflammasome assembly. (A–C) Differentiated *TRE-NLRP3-L353P*-THP-1 cells were loaded with 4 μM Fluo-8 for 1 h and treated with DOX (30 ng/mL) at 37°C for 6 h. The images were recorded by confocal microscopy at 5-min intervals from 3 h to 6 h. (A) Representative temporal subtraction images. (B) The frequency of intracellular Ca^2+^ increase at each time point. (C) The cumulative number of Fluo-8-positive cells. (D–F) Differentiated *TRE-NLRP3-L353P*-THP-1 cells were loaded with 4 μM Fluo-8 for 1 h and treated with DOX (30 ng/mL) at 37°C for 6 h in Ca^2+^-depleted or -supplemented media. The images were recorded by confocal microscopy at 5-min intervals from 3 h to 6 h. (D) Representative temporal subtraction images. (E) The frequency of intracellular Ca^2+^ increase at each time point. (F) The cumulative number of Fluo-8-positive cells. (G–I) Differentiated *EF1-ASC-BFP/TRE-NLRP3-L353P*-THP-1 cells were loaded with 4 μM Fluo-8 for 1 h and treated with DOX (30 ng/mL) at 37°C. The images were recorded at 5-min intervals. (G) Representative images of the cells with increased Fluo-8 signals. (H) The ASC-BFP speck size (I) and fluorescent intensity of Fluo-8 in were analyzed. The peak time point of Fluo-8 signals was defined as 0 min. (H) The blue line and the (I) green line represent mean values and the gray line represents each measurement. \**P* < 0.05, \*\**P* < 0.01, \*\*\**P* < 0.005 as determined by (C and F) Fisher’ s exact test with the Holm correction or (H and I) repeated one-way ANOVA with a post hoc test. (A–G) Data are representative of three independent experiments. (H and I) Data are from three independent live-cell imaging.

### Caspase-1 inhibition prevents mutated NLRP3-mediated inflammasome assembly

Recent studies have suggested that caspase-mediated pore formation by GSDMD causes the influx of Ca^2+^ (de Vasconcelos et al., 2019). We postulated that caspase-dependent Ca^2+^ influx might enhance inflammasome assembly. To investigate the effect of caspase-1 inhibition on Ca^2+^ influx, cells were treated with VX-765, a caspase-1 inhibitor, prior to DOX-mediated NLRP3-L353P induction. Indeed, VX-765 canceled Ca^2+^ influx (Figure 6A–C, Video 2, Figure 6–figure supplement 1A and B) and inhibited ASC oligomerization and speck formation induced by NLRP3-L353P (Figure 6D and E). In accordance with reduced inflammasome assembly, VX-765 also prevented NLRP3-L353P-induced IL-1β release (Figure 6F). These results indicate that caspase-1 activation induced by NLRP3 mutants promotes incremental inflammasome assembly by regulating Ca^2+^ influx. On the other hand, MCC950 (Coll et al., 2015), a potent NLRP3 inhibitor, failed to prevent ASC speck formation and IL-1β release induced by NLRP3-L353P, although MCC950 efficiently inhibited nigericin-induced IL-1β release (Figure 6–figure supplement 2A–C). These results suggest that caspase-1 inhibition could be a potential therapeutic target of CAPS.

**Figure 6.**
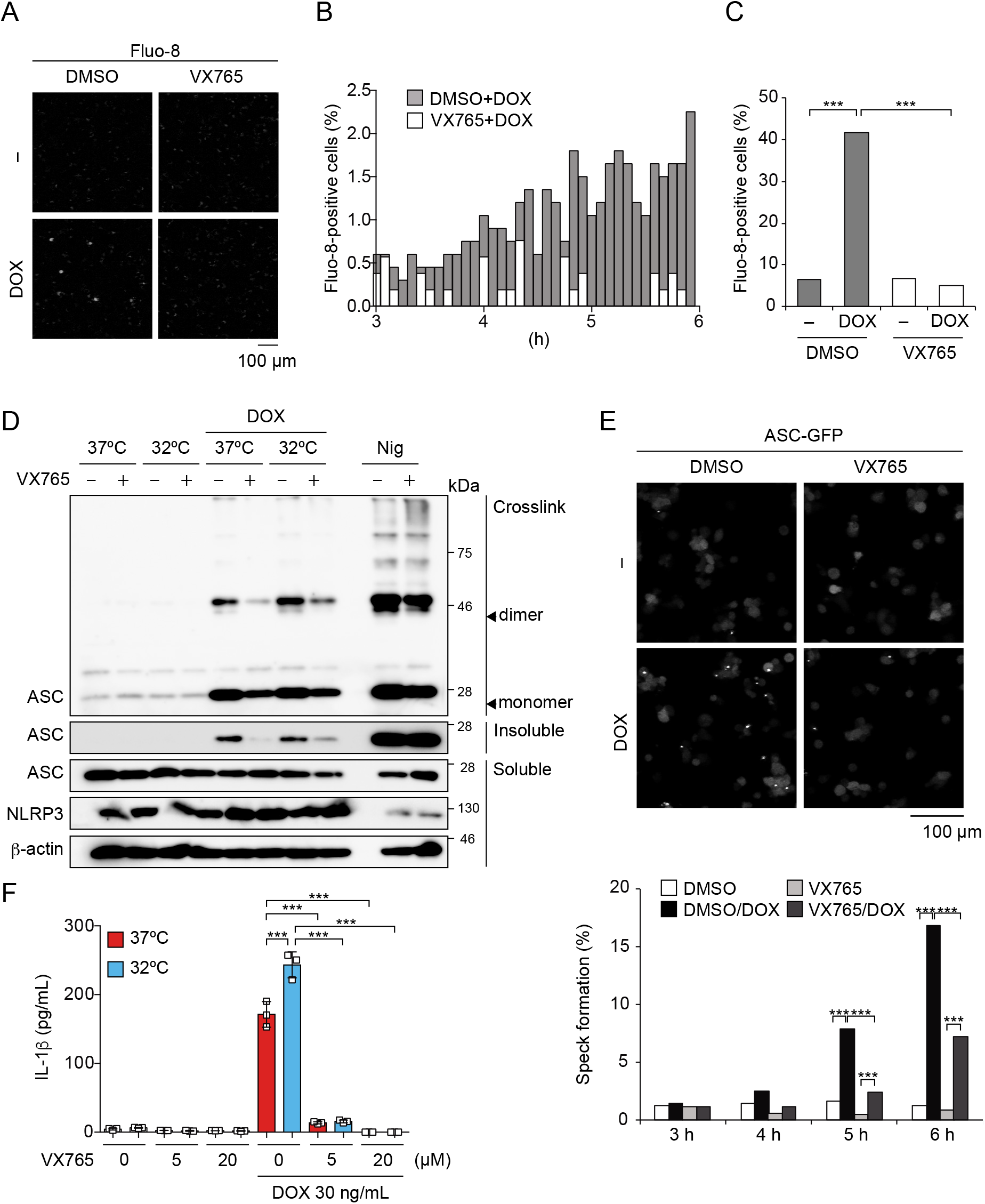
Caspase-1 inhibition prevents mutated NLRP3-mediated inflammasome assembly. (A–C) Differentiated *TRE-NLRP3-L353P*-THP-1 cells were loaded with 4 μM Fluo-8 for 1 h and were pretreated with VX-765 (20 μM) for 30 min. After DOX (30 ng/mL) treatment, the images were recorded at 5-min intervals from 3 h to 6 h. (A) Representative temporal subtraction images. (B) The frequency of intracellular Ca^2+^ increase at each time point. (C) The cumulative number of Fluo-8-positive cells. (D–F) Differentiated *TRE-NLRP3-L353P*-THP-1 cells were pretreated with VX-765 (20 μM) for 30 min and then treated with DOX (30 ng/mL) or nigericin (5 μM) at 37°C or 32°C. (D) Triton X-insoluble fractions were crosslinked with BS3 and analyzed by Western blot. (F) The IL-1β levels in the supernatants were assessed by ELISA (n = 3). (E) *EF1-ASC-GFP/TRE-NLRP3-L353P*-THP-1 cells were pretreated with 20 μM VX-765 for 30 min and then treated with DOX (30 ng/mL) at 37°C. ASC speck formation was analyzed by confocal microscopy. (F) Data are expressed as the mean ± SD. \**P* < 0.05, \*\**P* < 0.01, \*\*\**P* < 0.005 as determined by (C and E) Fisher’ s exact test with the Holm correction or (F) two-way ANOVA with a post hoc test. Data are representative of two or three independent experiments.

## Discussion

In the present study, we demonstrated that CAPS-associated NLRP3 mutants form cryo-sensitive foci intracellularly. These foci are aggregates that function as a scaffold for inflammasome activation. Consistent with this finding, inflammasome assembly and subsequent IL-1β release induced by CAPS-associated NLRP3 mutants are cryo-sensitive. The aggregation of CAPS-associated NLRP3 mutants is regulated by intracellular Ca^2+^ levels. Furthermore, caspase-1 inhibition prevents Ca^2+^ influx and inflammasome assembly induced by CAPS-associated NLRP3 mutants (Figure 7). These findings provide new insights into the molecular mechanisms of inflammasome activation in CAPS.

**Figure 7.**
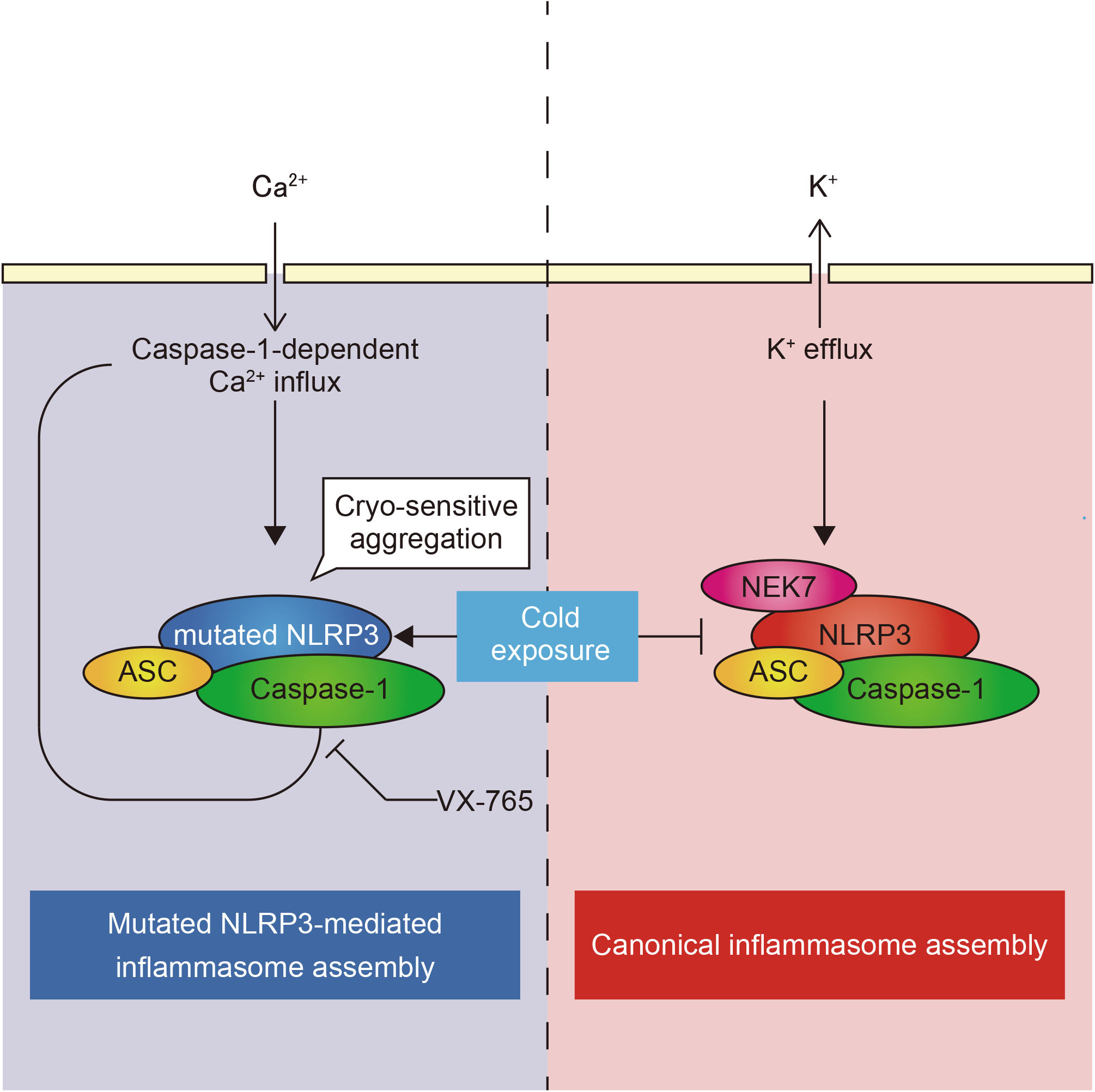
Model of CAPS-associated NLRP3 mutants-mediated inflammasome assembly. The mutated NLRP3 forms cryo-sensitive aggregates which trigger inflammasome assembly. The mutated NLRP3-mediated inflammasome assembly is sensitive to Ca^2+^ whereas canonical NLRP3 inflammasome assembly is dependent on K^+^ efflux. The inhibition of caspase-1 prevents the mutated NLRP3-mediated Ca^2+^ influx and inflammasome assembly. **Video 1. Caspase activity is required for Ca^2+^ influx induced by mutated NLRP3** Differentiated *TRE-NLRP3-L353P*-THP-1 cells were loaded with 4 μM Fluo-8 for 1 h and treated with DOX (30 ng/mL) at 37°C for 6 h in Ca^2+^-depleted or -supplemented media. The images were recorded by confocal microscopy at 5-min intervals from 3 h to 6 h.

**Figure 5 – figure supplement 1.**
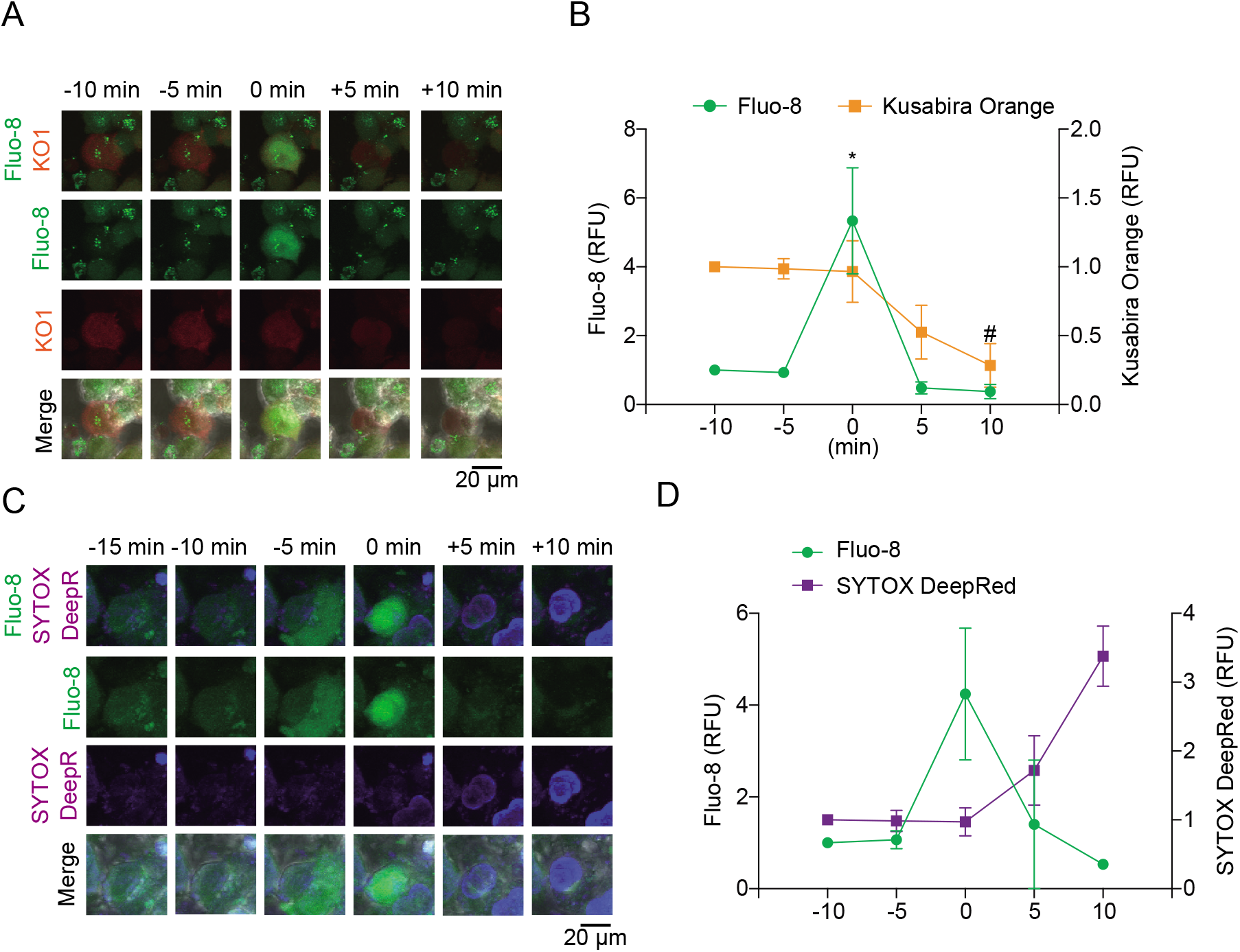
Pyroptosis is induced by mutated NLRP3 after Ca2+ influx. (A and B) Differentiated *TRE-NLRP3-L353P*-THP-1 cells were loaded with 4 μM Fluo-8 for 1 h and treated with doxycycline (30 ng/mL) at 37°C for 6 h. (A) Representative images of cells with increased Fluo-8 signal. (B) The relative fluorescent intensity of Fluo-8 and Kusabira Orange in cells with increased Fluo-8 signal was quantified (n = 10). (C and D) Differentiated *TRE-NLRP3-L353P*-THP-1 cells were loaded with 4 μM Fluo-8 for 1 h and treated with DOX (30 ng/mL) at 37°C for 6h in the presence of SYTOX DeepRed. (C) Representative images of cells with increased Fluo-8 signal. (D) The relative fluorescent intensity of Fluo-8 and SYTOX DeepRed in cells with increased Fluo-8 signal was quantified (n = 5). Data are expressed as the mean ± SD. \**P* < 0.05, as determined by repeated one-way ANOVA with a post hoc test. Data are analyzed from two independent time-lapse imaging.

**Figure 6 – figure supplement 1.**
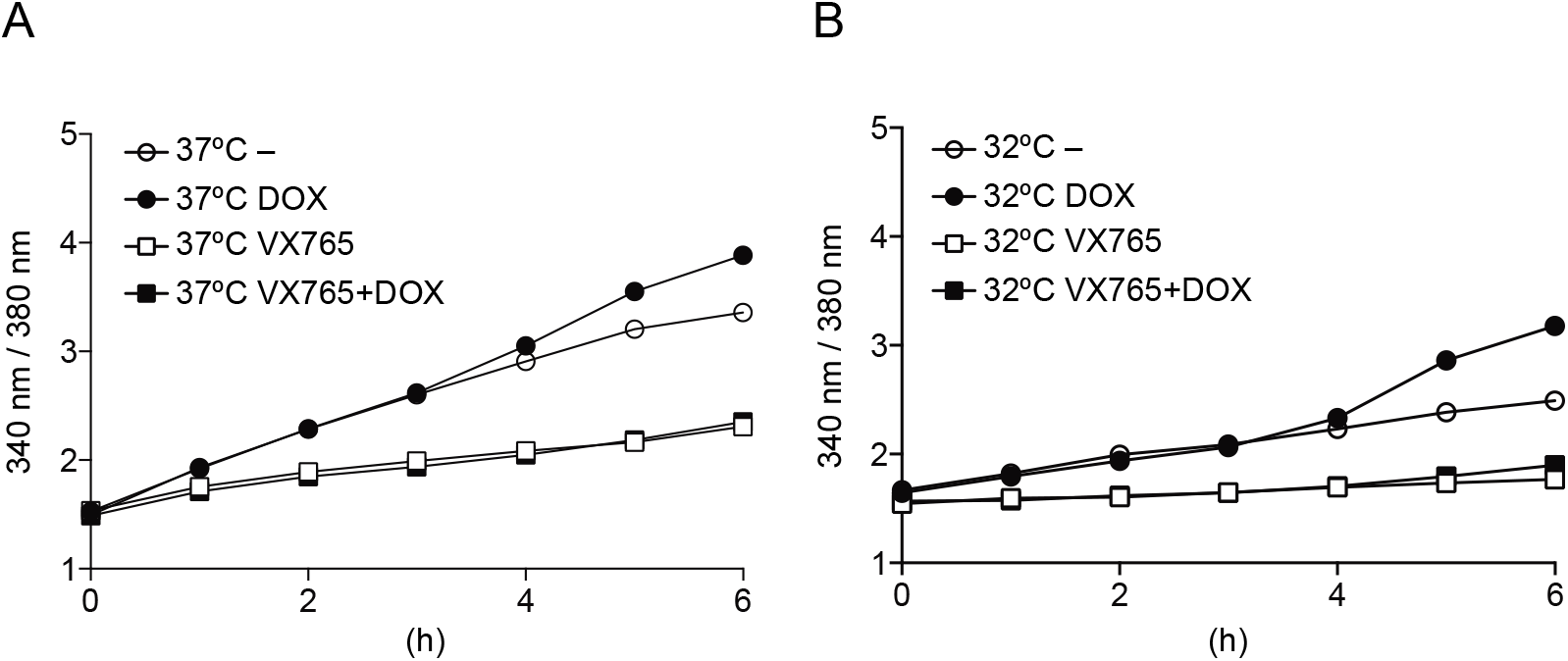
Caspase activity is required for Ca^2+^ influx induced by mutated NLRP3. (A and B) Differentiated *TRE-NLRP3-L353P*-THP-1 cells were loaded with 3 μM Fura2 for 1 h. Cells were pretreated 20 μM VX765 for 30 min and then treated with DOX (30 ng/mL) at 37°C or 32°C for 6 h. Fluorescence (A) at 37°C and (B) at 32°C was measured at hourly interval (n = 6). Data are expressed as the mean ± SD. Data are representative of two independent experiments. **Video 2. Caspase activity is required for Ca^2+^ influx induced by mutated NLRP3** Differentiated *TRE-NLRP3-L353P*-THP-1 cells were loaded with 4 μM Fluo-8 for 1 h and were pretreated with VX-765 (20 μM) for 30 min. After DOX (30 ng/mL) treatment, the images were recorded at 5-min intervals from 3 h to 6 h.

**Figure 6 – figure supplement 2.**
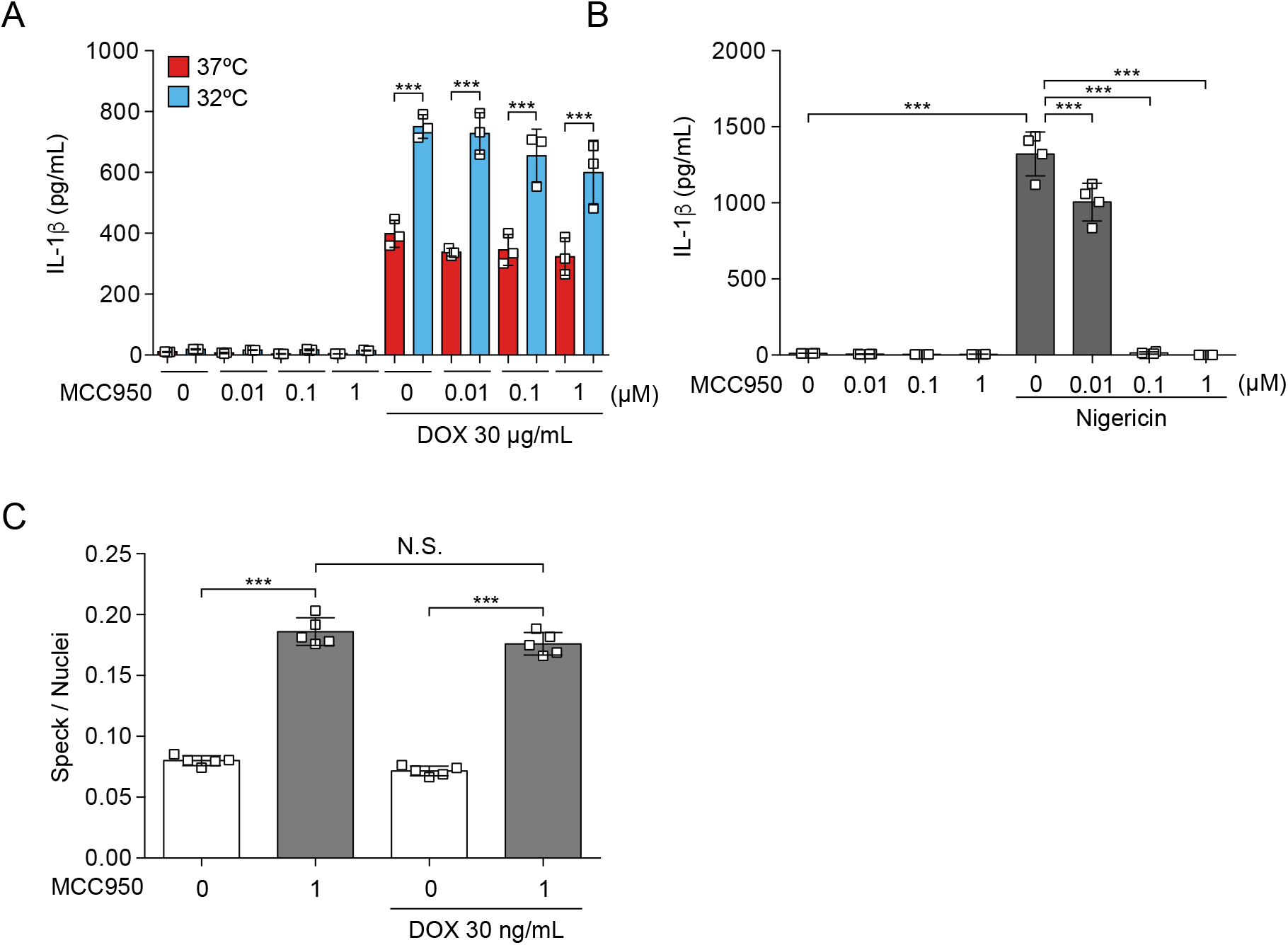
MCC950 failed to inhibit mutated NLRP3-induced inflammasome activation. (A–B) Differentiated *TRE-NLRP3-L353P*-THP-1 cells were pretreated with MCC950 for 30 min and then treated with (A) DOX (30 ng/mL) or (B) nigericin (5 μM) at 37°C or 32°C. The levels of IL-1β in the supernatants were determined by ELISA (n=3). (C) *EF1-ASC-GFP/TRE-NLRP3-L353P*-THP-1 cells were pretreated with MCC950 and then treated with DOX (30 ng/mL) at 37°C. The formation of ASC-speck was analyzed by high content analysis. Data are expressed as the mean ± SD. \*\*\**P* < 0.005 as determined by one-way ANOVA with a post hoc test. Data are representative of two independent experiments.

The formation of cryo-sensitive aggregates by CAPS-associated NLRP3 mutants is a significant finding of this study. Oligomerization or polymerization is a common feature of domains contained in inflammasome components. Both CARD in ASC and caspase-1 and PYD in ASC and NLRP3 form filamentous assemblies (Masumoto et al., 1999; Lu et al., 2014; Karasawa et al., 2015; Stutz et al., 2017). In addition, full-length ASC, which has both PYD and CARD, forms a large aggregate called speck. Therefore, aggregation of CAPS-associated NLRP3 mutants seems to be mediated by their PYD-PYD interaction. However, the mechanisms by which CAPS-associated NLRP3 mutants, which typically occur in other domains including NACHT domain and LRR, promote aggregation and affect their cryosensitivity have remained unclear. According to the structural analysis, an unstructured domain is present between PYD and NACHT. Although the structure of NLRP3 without PYD has recently been determined by cryo-electron microscopy, the structure of full-length NLRP3 and the effect of CAPS-associated mutation on it is unclear (Sharif et al., 2019). This information would be useful for clarifying how CAPS-associated NLRP3 mutants form aggregates.

With regard to cryo-sensitivity, recent studies have suggested that the formation of aggregates and LLPS is modulated by conditions including temperature and pH (Alberti et al., 2019; Riback et al., 2020). These factors shift the threshold of aggregation and LLPS under a constant protein concentration. In the present study, the frequency of foci formation by mutated NLRP3 was weakly associated with its expression levels. Furthermore, increased expression of mutated NLRP3 dose-dependently promoted inflammasome assembly. The temperature and its expression levels seem to be an essential determinant of the aggregation of mutated NLRP3.

Increasing evidence suggests that K^+^ efflux is a main upstream event of NLRP3 inflammasome activation (Munoz-Planillo et al., 2013; He et al., 2016a). Although the precise mechanism underlying K^+^ efflux-mediated NLRP3 inflammasome assembly is still unclear, NEK7 has been shown to interact with NLRP3 directly and promote inflammasome assembly as a downstream of K^+^ efflux (He et al., 2016b). Unexpectedly, we demonstrated that inflammasome activation induced by NLRP3-L353P is not prevented by supplementation with extracellular K^+^ or a deficiency of NEK7, indicating that K^+^ efflux is dispensable for inflammasome activation induced by FCAS-associated NLRP3 mutant. Instead, our data clearly showed that both NLRP3 aggregation and inflammasome assembly induced by NLRP3-L353P are dependent on the presence of Ca^2+^. Several studies indicate that Ca^2+^ is another upstream signal of NLRP3 inflammasome activation (Lee et al., 2012; Murakami et al., 2012; Rossol et al., 2012; Horng, 2014). Further studies are necessary to elucidate whether the previously identified mechanism of Ca^2+^ -mediated WT NLRP3 inflammasome activation shares the mechanism with inflammasome activation induced by CAPS-associated NLRP3 mutants. However, the Ca^2+^-sensitive NLRP3 aggregation appear to be distinct from canonical inflammasome assembly because the stimulation with nigericin failed to form NLRP3 aggregates.

Of note, we showed that the elevation of intracellular Ca^2+^ induced by NLRP3-L353P was dependent on caspase-1 activity. Recent studies have suggested that gasdermin pore formation induced by caspase activation regulates the influx of Ca^2+^ (de Vasconcelos et al., 2019). Broz and his colleagues have suggested that the Ca^2+^ influx induced by GSDMD pore formation initiates membrane repair by ESCRT complex, which negatively regulates pyroptosis (Rühl et al., 2018). Thus, the elevation of intracellular Ca^2+^ by GSDMD pore seems to regulate various cellular functions. In the present study, pharmacological caspase-1 inhibition prevents the elevation of intracellular Ca^2+^ and ASC oligomerization and speck formation. Therefore, we consider that CAPS-associated NLRP3 mutant forms cryo-sensitive inflammasome assemblies, which trigger caspase-1-mediated feed-forward loop of Ca^2+^ influx, leading to incremental inflammasome assembly. These findings also suggest that caspase-1 inhibition could be a potential target of inflammasome assembly induced by CAPS-associated NLRP3 mutants.

In the present study, we investigated the mechanisms of cryo-sensitive inflammasome assembly in CAPS-associated NLRP3 mutants. However, this study has a few limitations. First, to analyze the function of mutated NLRP3, we used mutated NLRP3 expressing under an inducible promoter or in *ASC KO* cells because the expression of mutated NLRP3 induces pyroptotic cell death. To clarify the dynamics of endogenously expressed NLRP3, a study using a knock-in model would be required. Second, the aggregation of NLRP3 was analyzed by fusion protein with mNeonGreen. A biochemical analysis of aggregation using mutated NLRP3 without any tags or fluorescent proteins is also required in the future.

In conclusion, we found that CAPS-associated NLRP3 mutants form cryo-sensitive aggregates, which can function as the scaffold for NLRP3 inflammasome activation. The aggregation of mutated NLRP3 is not dependent on K^+^ efflux but rather is regulated by intracellular Ca^2+^ levels. We expect that our findings would be valuable for the development of novel therapies for CAPS.

## Author Contributions

Conceptualization, T.Ka., and M.T.; Methodology, E.A., T.Ka., and S.W.; Validation, T.Ko., Y.M., and N.Y; Investigation, T.Ka., E.A., S.W., T.Ko., and C.B.; Resources, E.A., T.Ka; Writing-Original Draft, T.Ka.; Writing-Review & Editing, T.Ka., M.T.; Visualization, T.Ka.; Supervision, T.Ko., T.M., and M.T.; Project Administration, T.Ka., and M.T.; Funding Acquisition, T.Ka. and M.T.

## Acknowledgments

This study was supported by the Japan Society for the Promotion of Science (JSPS) through the Grants-in-Aid for Scientific Research (C), (18K08112, M.T.; 18K08485, T.Kar.), Grants-in-Aid for Scientific Research on Innovative Areas (Thermal Biology) (16H01395. M.T.), the Agency for Medical Research and Development-Core Research for Evolutional Science and Technology (AMED-CREST) (M.T.), and the Ministry of Education, Culture, Sports, Science and Technology (MEXT)-supported program for Private University Research Branding Project (M.T.). We are grateful to Dr. Kunitoshi Uchida and Dr. Jun Fujita for their valuable suggestions. We thank Naoko Sugaya, Masako Sakurai, and Rumiko Ochiai for their technical assistance.

## Abbreviations

ASC: apoptosis-associated speck-like protein containing a caspase recruitment domain
CAPS: cryopyrin-associated periodic syndromes
CINCA: chronic infantile neurological, cutaneous and articular syndrome
DAMP: damage/danger-associated molecular pattern
DOX: doxycycline
FCAS: familial cold autoinflammatory syndrome
FRAP: fluorescence recovery after photobleaching
GSDMD: gasdermin D
IL: interleukin
LLPS: liquid-liquid-phase separation
MWS: Muckle-Wells syndrome
NLRP3: nucleotide-binding oligomerization domain, leucine-rich repeat and pyrin domain containing 3
PRR: pattern recognition receptor
RCD: regulated cell death

## METHODS

### KEY RESOURCES TABLE

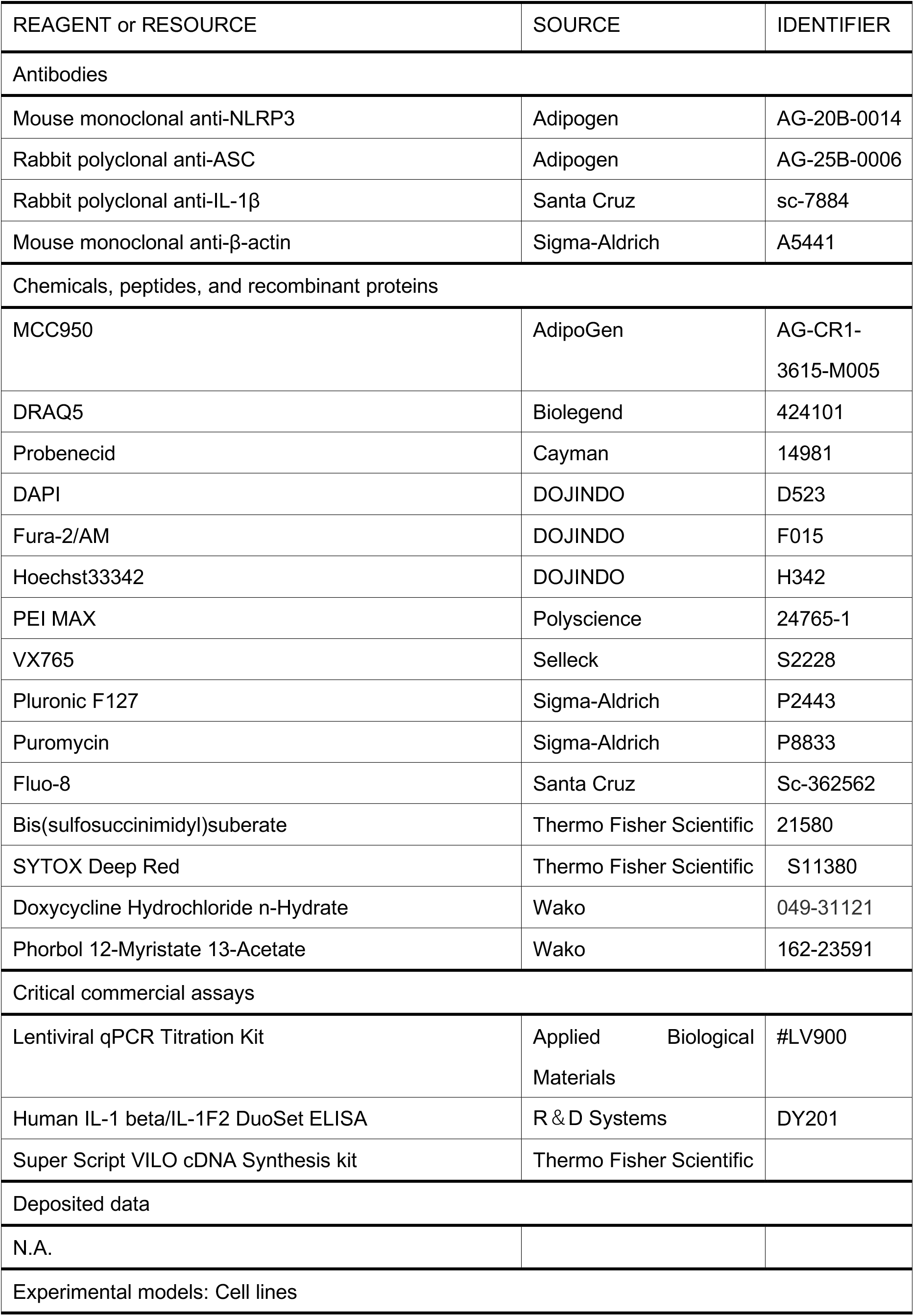
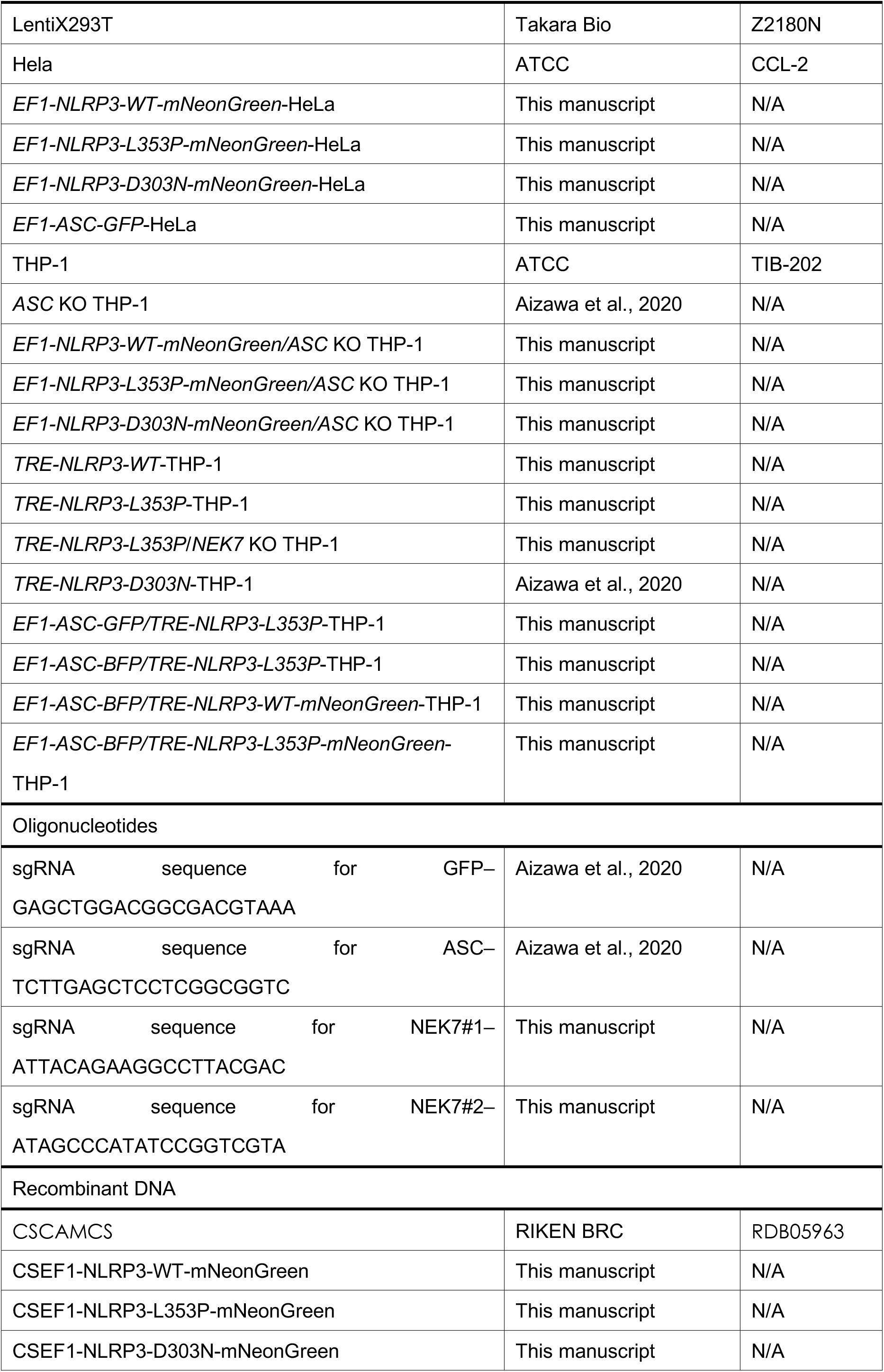
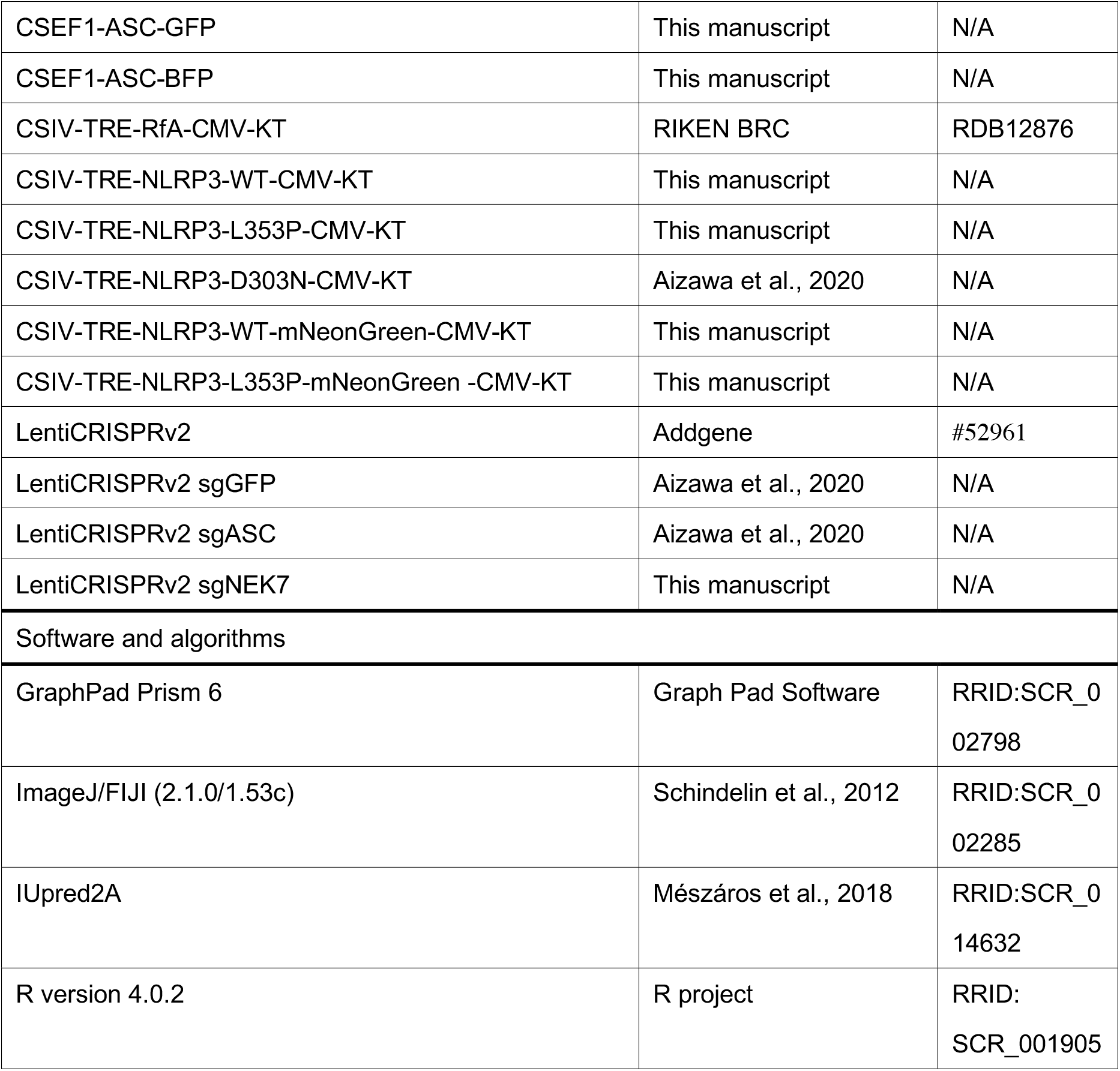

### Plasmids

Polymerase chain reaction (PCR)-generated cDNAs encoding human NLRP3 were subcloned into pENTR4 vector. The mutated NLRP3-D303N and NLRP3-L353P were generated using the PrimeSTAR Mutagenesis Basal kit (Takara Bio, Shiga, Japan) (Aizawa et al., 2020). The primers for introducing mutations were as follows: D303N (forward, 5′-GGCTTCAATGAGCTGCAAGGTGCCTTTGACGAG-3′; reverse, 5′-CAGCTCATTGAAGCCGTCCATGAGGAAGAGGAT-3′), L353P (forward, GTGGCCCCGGAGAAACTGCAGCACTTGCTGGAC-3′; reverse, 5′-TTTCTCCGGGGCCACAGGTCTCGTGGTGATGAG-3′). To produce DOX-inducible expression vector, the mutated NLRP3 and WT NLRP3 were transferred into CS-IV TRE CMV KT (kindly provided by Dr. H. Miyoshi, RDB12876, RIKEN BRC, Tsukuba, Japan) with LR clonase (Thermo Fisher Scientific). To develop NLRP3-mNeonGreen or ASC-BFP-expressing lentiviral vector, PCR-generated NLRP3-WT, NLRP3-L353P, NLRP3-D303N, ASC, mNeonGreen, moxBFP (a gift from Erik Snapp; Addgene plasmid #68064) were Gibson subcloned into CS-EF-1 (derived from CS-CA-MCS; RIKEN BRC). The sgRNA targeting NEK7 was designed with CRISPR direct (http://crispr.dbcls.jp) and subcloned into LentiCRISPRv2, which was a gift from Feng Zhang (Addgene plasmid #52961; http://n2t.net/addgene: 52961; RRID: Addgene_52961).

### Cell culture

HeLa cells were cultured in Dulbecco’s modified Eagle’s medium (DMEM, Wako, Osaka, Japan) supplemented with 10% fetal calf serum (FCS) and antibiotics. THP-1 cells were cultured in RPMI1640 (Sigma, St Louis, MO, USA) supplemented with 10% fetal calf serum (FCS) and antibiotics. THP-1 macrophages were differentiated with 200 nM phorbol-12-myristate-13-acetate (PMA) for 24 h. LentiX293T cells were obtained from TAKARA (Takara Bio, Shiga, Japan) and cultured in DMEM supplemented with 10% FCS, 1 mM sodium pyruvate, and antibiotics. Unless otherwise indicated, cells were cultured at 37°C in 5% CO_2_.

### Lentiviral preparation

LentiX293T cells were co-transfected with self-inactivating vectors, pLP1, pLP2, and pVSVG using PEI MAX (Polysciences, Warrington, PA, USA) to prepare the lentiviral vectors. Culture media containing the lentiviral vectors were collected 3 days after transfection. The collected media were filtered with a 0.45-μm filter and ultracentrifuged at 21,000 rpm using a Type 45 Ti rotor (Beckman Coulter, Brea, CA, USA), and the pellets were resuspended in phosphate-buffered saline (PBS) containing 5% FCS. The lentivirus titer was measured using a Lentivirus qPCR Titer kit (Applied Biological Materials, Richmond, BC, Canada). For lentiviral transduction, the cells were incubated with purified lentiviral vectors in the presence of 8 μg/mL polybrene (Sigma). The details of the developed cells are described in the key resource table.

### Treatment of reagents and cold exposure

The transduced THP-1 cells were differentiated with 200 nM PMA for 24 h and treated with DOX (Wako) or nigericin (InvivoGen) at the indicated concentrations. Next, cells were cultured at 37°C or 32°C. Cells were then pretreated with inhibitors including CA-074 (Wako), MCC950 (Adipogen, San Diego, CA), VX-765 (Selleck), and Z-VAD-FMK (MBL) for 30 min prior to cold exposure.

### Confocal microscopy

For imaging of fixed cells, the transduced cells were seeded on an 8-well chamber slide (Matsunami Glass Ind., Ltd., Osaka, Japan), and then fixed with neutral buffered formalin or 1% paraformaldehyde and stained with 1 μg/mL 4’,6-diamidino-2-phenylindole, dihydrochloride (DAPI; Dojindo). For live-cell imaging, cells were seeded at 1×10^5^ cells/well on 8-well cover glass chambers (IWAKI, Shizuoka, Japan) and labeled with Hoechst33342 for 20 min before treatment. The images were captured using confocal microscopy (FLUOVIEW FV10i; Olympus, Tokyo, Japan).

### High content analysis

The transduced HeLa cells or THP-1 cells were seeded on 96-well plates and treated with the indicated stimuli. For analysis of fixed cells, nuclei were stained with DAPI after fixation by 1% paraformaldehyde. For live-cell imaging, cells were stained with DRAQ5 (Biolegend, San Diego, CA) and analyzed by an Operetta CLS high-content analysis system (PerkinElmer, Waltham, MA).

### FRAP analysis

The transduced HeLa cells were seeded on 8-well cover glass chambers (IWAKI), cultured at 32°C for 24 h, and analyzed by FV1000 confocal microscopy (Olympus) at 32°C. Images were captured using an UPLA SAPO 60XO objective. The GFP and mNeonGreen signals were captured using a line sequential scan setting with excitation laser lines at 488 nm. For FRAP analysis, a 1 sec pulse of the 488 nm laser line at 5% power was used to bleach the NLRP3-mNeonGreen foci. A 5 sec pulse of 488 laser line at 30% power was used to bleach the ASC-GFP specks. The changing of fluorescence was monitored by imaging every 0.2 msec.

### IL-1β secretion assay

Cells were seeded into 96-well plates at 5 ×10^4^ cells/well. After the indicated treatments, culture supernatants were collected and the IL-1β levels were measured by enzyme-linked immunosorbent assay (ELISA) using a commercial kit (R&D Systems, Minneapolis, MN, USA). The supernatants were precipitated with ice-cold acetone and resolved in 1 × Laemmli buffer for Western blot analysis.

### Western blot analysis

Samples were separated by sodium dodecyl sulfate-polyacrylamide electrophoresis (SDS-PAGE) and transferred to PVDF membranes. After blocking with Blocking One (Nacalai Tesque, Kyoto, Japan) for 30 min, the membranes were incubated overnight at 4°C with the following primary antibodies (Abs): anti-ASC (AL-177; Adipogen), anti-β actin (clone AC-15; Sigma), anti-Caspase-1 (3866; Cell Signaling Technology), anti-IL-1β (H153; Santa Cruz Biotechnology), and anti-NLRP3 (clone Cryo-2; Adipogen). As secondary Abs, HRP-goat antimouse Superclonal IgG (Thermo Fisher Scientific) or HRP-goat anti-rabbit IgG (Cell Signaling Technology) was incubated with membrane for 1 h. After being washed with TBS-Tween, immunoreactive bands were visualized using Western BLoT Quant HRP Substrate (Takara Bio) or Western BLoT Ultra Sensitive HRP Substrate (Takara Bio).

### ASC-oligomerization assay

Cells were lysed in 0.5% Triton X-100 buffer (20 mM Tris HCl, 10 mM KCl, 1.5 mM MgCl_2_, 1 mM EDTA, 1 mM EGTA, 320 mM sucrose, and 0.5% Triton X-100) for 20 min. Lysates were then centrifuged at 5000 xg for 10 min. The insoluble pellets were reacted with 2 mM bis(sulfosuccinimidyl)suberate (BS3) (Thermo Fisher Scientific) for 30 min and the reactions were terminated by an excess amount of glycine.

### Reverse transcription and real-time PCR

Total RNA was prepared using ISOGEN (Nippon Gene Co., Tokyo, Japan) according to the manufacturer’s instructions. Total RNA was reverse-transcribed using a SuperScript VILO cDNA Synthesis kit (Life Technologies). Real-time PCR was performed using SYBR Premix Ex Taq II (Takara Bio). The primers used in the assay were as follows: NEK7 (forward, 5′-GCCTTACGACCGGATATGGG-3′; reverse, 5 ‘-CACTAAATTGTCCGCGACCAA–3′), and ACTB (forward, 5′-GGCACTCTTCCAGCCTTCCTTC-3′; reverse, 5′-GCGGATGTCCACGTCACACTTCA-3′).

### Ca^2+^ -imaging using Fluo-8

Cells were seeded at 1×10^5^ cells/well on an 8-well cover glass chamber (IWAKI). Fluo-8 (Santa Cruz Biotechnology) was loaded at 4 μM for 60 min in the presence of 0.04% Pluronic F127 (Sigma)and 1.25 mM probenecid (Cayman). After removal of the loading medium, cells were treated with DOX in the presence of 1.25 mM probenecid. Z-stack time-lapse images at 37°C were captured using confocal microscopy (FLUOVIEW FV10). To normalize the cell number, nuclei were labeled with Hoechst 33342 (1 μg/mL) or DRAQ5. For detection of dying cells, images were captured in the presence of 100 nM SYTOX Deep Red.

### Fura2 Assay

Cells were seeded at 5×10^4^ cells/well into 96 well plates, and loaded with 3 μM fura 2-AM (Dojindo, Kumamoto, Japan) for 30 min. After cells were treated with the indicated reagents, fluorescence intensity (Excitation:340 or 380, Emission:510 nm) was measured by using a multimode microplate reader (Spark; TECAN, Switzerland) at 37°C or 32°C.

### Statistical analysis

Data are expressed as mean ± standard deviation (SD). Differences between two groups were determined by Student’s t-test. Differences between multiple group means were determined by two-way analysis of variance (ANOVA) combined with the Tukey’s post hoc test. Differences between multiple groups with repeated measurements were evaluated by repeated one-way ANOVA or repeated two-way ANOVA combined with the post hoc test. Analyses were performed using GraphPad Prism 6 software (Graph Pad Software, La Jolla, CA, USA) or R version 4.0.2 (https://www.r-project.org). A p-value of < 0.05 was considered statistically significant. Biological replicates indicate replicates of the same experiment conducted upon separately seeded culture on separate days. The number of biological replicates is described in the figure legend. For plate reader-based assay, n represents replicates that were acquired from different cells. In live-cell imaging assay, n represents replicates that were acquired from each cell through multiple set of experiments.

